# Uncovering the bequeathing potential of Apoptotic Mesenchymal Stem Cells via small Extracellular Vesicles for its enhanced immunomodulatory ability

**DOI:** 10.1101/2024.04.22.590581

**Authors:** Meenakshi Mendiratta, Mohini Mendiratta, Yashvi Sharma, Ranjit K. Sahoo, Neena Malhotra, Sujata Mohanty

## Abstract

Small Extracellular Vesicles (sEVs) derived from Mesenchymal Stem Cells (MSCs) have emerged as a promising avenue for cell-free therapeutics in regenerative medicine. These vesicles, endowed with regenerative cargo inherited from their parent cells, have attracted attention for their role in immunomodulation and ROS alleviation. Notably, the deliberate induction of apoptosis in MSCs prior to Extracellular Vesicles (EVs) isolation has been identified as a strategy to augment the regenerative capabilities of MSCs-EVs, as certain reports have suggested that MSCs undergo apoptosis to exert their therapeutic effect post-transplantation. Moreover, selecting an optimal tissue source for deriving MSC-sEVs is equally crucial to ensure consistent and improved clinical outcomes.

Multiple attributes of MSCs like their antioxidant, Immunomodulatory & regenerative properties make them particularly appealing for clinical studies, wherein mechanisms such as paracrine secretions and efferocytosis play pivotal roles. This investigation meticulously explores the comparative immunomodulatory & antioxidant capabilities of Apoptotic sEVs (Apo-sEVs) with Viable sEVs (V-sEVs) obtained from both Bone Marrow (BM) and Wharton’s Jelly (WJ)-derived MSCs, using an *in vitro* liver injury model. The findings from the present study contribute valuable insights into the comparative efficacy of Apo-sEVs and V-sEVs. This will aid in addressing a critical gap in understanding the role of apoptosis in enhancing the reparative capability of MSCs-sEVs. It also aims to shed light on the optimal source of MSCs for generating Apo-sEVs in translational applications.

**GRAPHICAL ABSTARCT:** 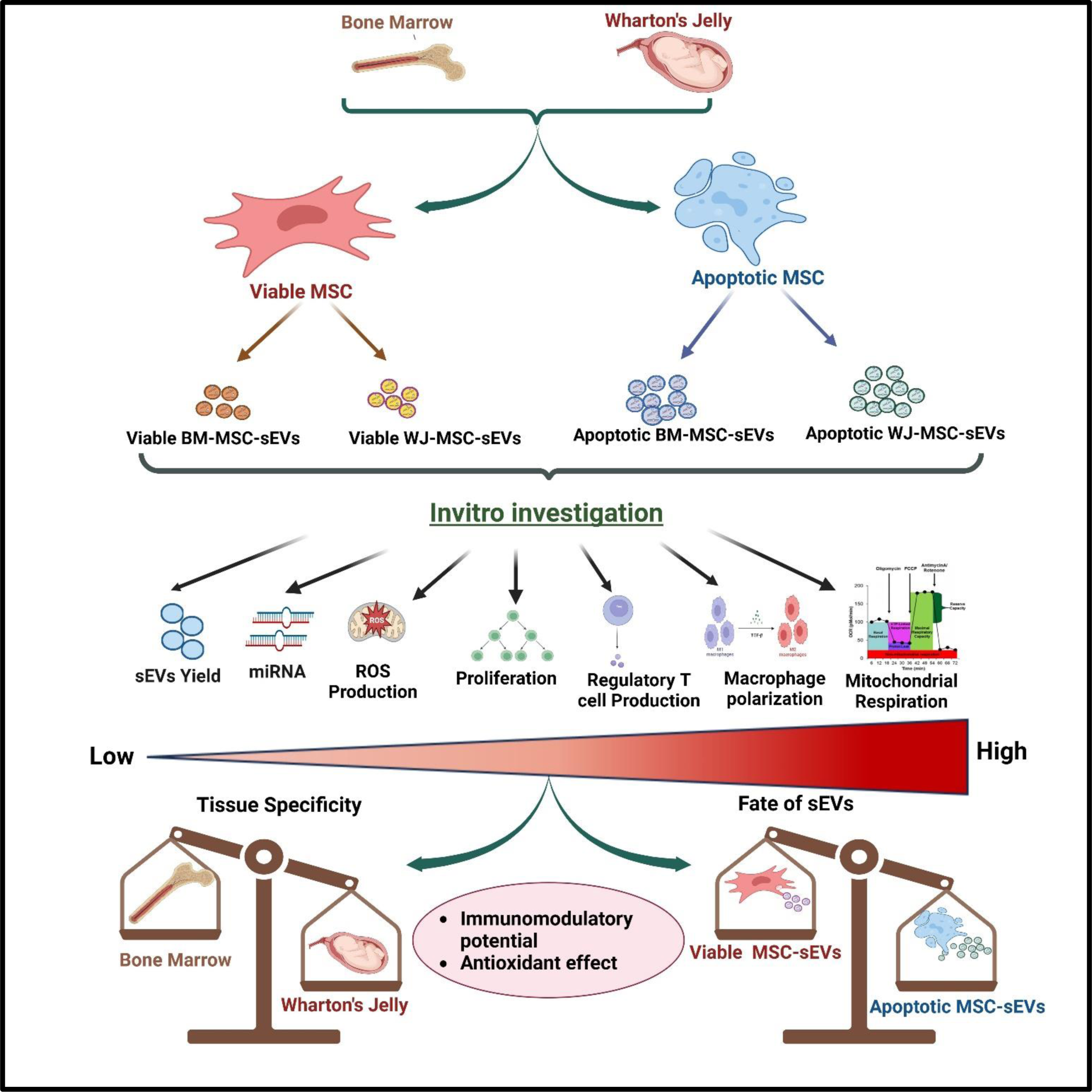

## Introduction

Mesenchymal stem cells (MSCs) are multipotent stem cells, that can be isolated from various adult tissues, including bone marrow, umbilical cord, adipose tissue, and dental pulp[1,2]. The attainment of effective clinical outcomes relies on the utilization of MSCs from an optimal tissue source, which can consistently yield improved clinical results, along with posing least ethical concerns, being easily available & having cost efficacy [3]. MSCs have garnered significant attention in clinical trials due to their regenerative, anti-oxidant, and immunomodulatory properties [4,5]. These properties are attributed to several mechanisms of action exhibited by MSCs, predominantly including paracrine secretions & efferocytosis by the act of apoptosis [6,7].

A key modality that has come to light in the context of paracrine secretions by MSCs is the sEVs secreted by them. These vesicles are less than 200 nm in size & are majorly involved in intercellular communication. They carry regenerative cargo including miRNAs, mRNAs, and proteins, but are devoid of nuclei & therefore safer for transplantation as compared to their parent cells [8]. Additionally, sEVs are versatile for diverse formulations and provide convenient storage and transportability[9,10].

Certain reports have also suggested that transplanted MSCs have a limited lifespan in recipients. They undergo apoptosis either within the host circulation or in the engrafted tissue as a means of getting efferocytosed thereby disseminating their regenerative components [11]. Therefore, this regeneration mechanism has been explored in a few studies by deliberately inducing apoptosis in the MSCs & then validating their regenerative potential [11]. These Apoptotic MSCs (Apo-MSCs) have been found to possess the ability to induce immunosuppressive effects in animal models of lung injury, liver injury, and spinal cord injury [12,13]. Pang et al. (2021) in their study, inhibited the apoptotic induction in MSCs by silencing the apoptotic effectors BAK & BAX. They observed that there was a reduction in their immunomodulatory capabilities in a viable form, when administered in an asthmatic mice model, thereby potentiating the need of MSCs for undergoing apoptosis to exert their functional effect [11,14].

However, employing whole cells for infusion comes with limitations, including the risk of adverse reactions due to the heterogeneity of apoptotic cells, which may also encompass necrotic cells [15,16]. Consequently, a cell-free approach utilizing Apoptotic Extracellular Vesicles (Apo-EVs), including both small & large EVs, has been explored for its potential to mimic therapeutic effects similar to their parent cells [17]. The potential of Apo-EVs has been explored in multiple studies [18–20]. Zheng et al., (2021) observed in their study that Apoptotic Vesicles released by MSCs were able to confer functional modulations to liver macrophages via efferocytosis, and mitigate type 2 diabetes mellitus [21]. This presents a novel insight that Apo-EVs could be a functionally potent modality for targeting immune implicated liver diseases.

While majority of studies with Viable MSCs (V-MSCs) have confirmed the superior regenerative potential of small EVs, as compared to the larger EVs, existing studies with Apo-MSCs have considered Apo-EVs as a collective heterogenous population of small EVs, large EVs & Apoptotic bodies, which may confer pro- or anti-therapeutic effects due to the packaging of detrimental components synthesised as a part of the cell death processs [18–20]. To the best of our knowledge, the comparative functional capabilities of V-MSCs-sEVs and Apo-MSCs-sEVs is yet to be elucidated. Furthermore, there remains a gap in understanding the optimal source of MSCs for the generation of Apo-sEVs in the context of translational applications [19,20]

Hence, in this investigation, we have thoroughly examined Apoptotic sEVs (Apo-sEVs) in comparison to Viable sEVs (V-sEVs) obtained from both Bone Marrow (BM) and Wharton’s Jelly (WJ)-derived MSCs. The objective was to unveil their immunomodulatory and antioxidant capabilities in an *in vitro* liver injury model while evaluating apoptosis as a priming strategy to amplify the functional potential of MSCs-sEVs.

## Materials and Methods

### 1. Reagents and antibodies

High Glucose (HG)-Dulbecco’s modified Eagle medium (DMEM) and Low Glucose (LG)-DMEM were purchased from Gibco, USA. Fetal Bovine Serum (FBS) was purchased from Himedia, India. STEMPRO® MSC Serum Free Medium CTS, 100X Antibiotic-Antimycotic solution, dead cell apoptosis kit with Annexin V, FxCycle™ PI/RNase Staining Solution kit, Pierce™ BCA Protein Assay Kit were purchased from Thermo Fischer Scientific, USA. Staurosporine, PKH26 Red Fluorescent cell linker kit, Cell Counting Kit-8 kit, pH Rodo Red succinidyl ester, phorbol 12-myristate 13-acetate (PMA), and radioimmunoprecipitation assay lysis buffer was purchased from Sigma, USA. CD-63, ALIX, Flotillin, calnexin, and GAPDH were purchased from Genetex, USA and cleaved caspase-3 was purchased from CST, USA). CD206, Arginase, and iNOS were purchased from eBiosciences, USA. Cell TraceTM CFSE cell proliferation assay kit was purchased from BD Biosciences, US. MitoSOX red was purchased from Life Technologies, USA. Qiazol, PCR starter kit, and miRNeasy mini kit were purchased from Qiagen, Netherlands. SYBR Green Master Mix and Protease inhibitors were purchased from Promega, US. CD105, CD90, CD73, CD29, HLA-class I, HLA-class II, and CD34/45 (BD Pharmingen, France). Phalloidin and TRIzol reagent was purchased from Invitrogen, United States. High-Capacity cDNA Reverse Transcription Kit was purchased from Applied Biosystems, Invitrogen, United States.

### 2. Cell lines

HUH7 cells were maintained in HG-DMEM supplemented with 10% FBS and 1% Antibiotic-Antimycotic solution. THP 1 and Jurkat cells were maintained in RPMI-1640 medium supplemented with 10% FBS and 1% Antibiotic-Antimycotic solution.

### 3. Isolation and characterization of human MSCs

MSCs used in this study were isolated from healthy donors with their written consent after obtaining ethical clearance from the Institutional Committee for Stem Cell Research (IC-SCR) (Ref No. IC-SCR/140/23 (o)), AIIMS, New Delhi. MSCs obtained from the bone marrow (BM-MSCs), and Wharton’s Jelly (WJ-MSCs) were used in this study. The bone marrow was collected from the patient’s donor (n=3) who was undergoing the routine medical test procedure at Dr. B. R. Ambedkar IRCH, AIIMS, New Delhi. Briefly, BM-MSCs were isolated and cultured in LG DMEM containing 10% MSC grade FBS, and 1% Antibiotic-Antimycotic solution at 37°C in a 5% CO_2_ incubator. Umbilical cord (n=3) was collected in transport media and processed within 24h of normal or cesarean delivery of the donors with the age group 20-35 years from the Department of Obstetrics and Gynecology, AIIMS, New Delhi. Briefly, the umbilical cord was collected and washed with 1X PBS containing 1% Antibiotics. The cord was cut to expose Wharton’s Jelly part, chopped into small pieces using a surgical blade. The Wharton’s Jelly was placed in 35mm culture dish containing LG-DMEM with 10% MSC grade FBS and 1% Antibiotic-Antimycotic solution and incubated overnight at 37°C and 5% CO_2._ MSCs were sub-cultured at 70% confluency and characterized using Flow Cytometry and Trilineage Differentiation. Surface markers CD105, CD90, CD73, CD29, HLA-class I, HLA-class II, and CD34/45 were studied using Flow Cytometry (Beckman Coulter, USA) and analyzed using Kaluza Software Version 2.1. Tri-lineage differentiation (osteocyte, adipocyte, and chondrocyte) was performed as per previously established protocol [1,22].

To achieve homogeneity, we pooled BM-MSCs (n=3) and WJ-MSCs (n=3) at passage 3 for subsequent experiments.

### 4. Induction of apoptosis in MSCs

Both BM-MSCs and WJ-MSCs were treated with 0.5μM staurosporine (STS) in LG-DMEM for 6, 12 and 24 hours [19,20] The respective cell suspensions were collected, washed, and stained with Annexin V and PI to evaluate the apoptosis rate of apoptotic cells by flow cytometry and analyzed using Kaluza Software Version 2.1 and confirmed by cleaved caspase 3 expression via western blotting.

### 5. Isolation and characterization of Apoptotic MSC-derived small Extracellular Vesicles (Apo-sEVs)

Apo-sEVs were isolated from Apoptotic BM-MSCs (Apo-BM-MSCs) and Apoptotic WJ-MSCs (Apo-WJ-MSCs) using a one-step sucrose ultracentrifugation-based method by Optima XPN100 ultracentrifuge in a swinging bucket rotor (Beckman Coulter, USA) [22,23]. Briefly, Apo-MSCs were cultured in STEMPRO® MSC Serum Free Medium CTS for 48 hours, and the conditioned medium was collected for the isolation of Apo-sEVs. The conditioned medium was centrifuged at 300×g for 10 minutes to remove cellular debris followed by 10,000×g for 30 minutes to remove microvesicles. The supernatant was collected without disturbing the pellet and added over 30% sucrose solution layer. Then, it was centrifuged at 100,000×g at 4°C for 90 minutes. The sucrose layer was collected and washed with phosphate-buffered saline (PBS) at 100,000 × g for 90 minutes to obtain Apo-sEVs pellets. Apo-sEVs were resuspended in 500 µL PBS and stored at −80 °C until further use. V-sEVs were isolated from BM-MSCs and WJ-MSCs as per the above-described protocol. V-BM-sEVs and V-WJ-sEVs were used as a control.

Apo-sEVs and V-sEVs were quantified using a Bicinchoninic Acid (BCA) assay (Pierce™ BCA Protein Assay Kit)

### 6. Characterization of sEVs

The V-sEVs and Apo-sEVs were characterized as per the MISEV Guideline with the following techniques:

#### a. Nanoparticle tracking analysis

The sEVs were diluted (1:10) in PBS and acquired for particle concentration and size distribution using NanoSight LM20 (Malvern Instruments Company, United Kingdom). The Brownian motion of each particle was tracked between frames and the size was calculated using the Stokes-Einstein equation [22].

#### b. Zeta potential

The sEVs were acquired for charge distribution using Malvern Nano zeta sizer (Malvern Instruments Company, United Kingdom). The zeta potential of each particle was tracked between frames.

#### c. Transmission electron microscopy (TEM)

The sEVs were diluted at a concentration of 1:10 in PBS and placed on Formvar-carbon-coated copper grids and adsorbed for 5 minutes. The grids were stained with 2% Phosphotungstic acid solution for 1 minute. Then, grids were air-dried and observed under the transmission electron microscope (Tecnai, FEI, USA) [22].

#### d. Western blotting

The sEVs were lysed using radioimmunoprecipitation assay lysis buffer with protease inhibitors. Concentration of sEV lysates was estimated using a BCA protein assay kit. 25μg of sEVs lysate was subjected to 12% SDS-PAGE gel in reducing conditions for ALIX, calnexin, Cleaved Caspase3 and non-reducing condition for CD-63. This was followed by transfer to a PVDF membrane (MDI) using a wet transfer system (Bio-Rad, USA). The blot was blocked with 5% BSA in 1×TBS-T solution at room temperature for 1 hour followed by incubation with respective primary antibody CD-63 (1:500 dilution), ALIX (1:1000 dilution), calnexin (1:3000 dilution), cleaved caspase 3 (1:1000 dilution) and GAPDH (1:3000 dilution) overnight at 4°C. The blot was then washed with 1X TBST three times each for five minutes and incubated with HRP-conjugated secondary antibody for 2hours at room temperature. The blot was washed three times using 1XTBS-T and developed using an ECL imager (Bio-Rad, USA) [22].

### 7. Macrophage polarization

To evaluate the immunomodulatory potential of Apo-sEVs, THP-1 cells were seeded at a density of 1×10^6^ cells/well in a 6-well plate. THP-1 cells were differentiated into M0-like macrophages using 25 ng/ml PMA for 24 h in RPMI-1640 medium. M0-like macrophages were then polarized into M1-macrophages by treatment with 10 ng/ml of Lipopolysaccharide (LPS) for 24 h [24]. M1-macrophages were treated with 50 µg/mL V-BM-sEVs, V-WJ-sEVs, Apo-BM-sEVs, and Apo-WJ-sEVs for 48 hours. The treated macrophages were subsequently characterized with CD206-PE, Arginase 1-APC, and iNOS-FITC antibodies using flow cytometry and analyzed using Kaluza Software Version 2.1.

### 8. *In vitro* efferocytosis assay

In order to study the uptake of Apo-sEVs by the macrophages, we incubated the M1-macrophages with 20ng/ml pH Rodo Red succinidyl ester labeled V-BM-sEVs, V-WJ-sEVs, Apo-BM-sEVs, and Apo-WJ-sEVs [11]. Concurrently, M1-macrophages were labeled with 1µM CFSE dye for 20 minutes. The CFSE stained M1-macrophages were co-cultured with labeled V-BM-sEVs, V-WJsEVs, Apo-BM-sEVs, and Apo-WJ-sEVs for 48h. The percentage of Apo-sEVs engulfed by M1 macrophages was measured using flow cytometry and analyzed using Kaluza Software Version 2.1.

### 9. T-cell proliferation and differentiation assay

Jurkat cells were labeled with 1µM CFSE dye for 20 minutes followed by stimulation with PHA (5µg/ml) and IL-2 (50IU/ml) for 48hr. Afterward, stimulated Jurkat cells were treated with 50 µg/mL V-BM-sEVs, V-WJ-sEVs, Apo-BM-sEVs, and Apo-WJ-sEVs for 3 days. Furthermore, the proliferation and activation capacity of Jurkat cells were examined by Cell TraceTM CFSE cell proliferation assay kit and fluorochrome-conjugated antibodies such as CD3, CD4 respectively using a flow cytometer (Beckmann Coulter, US). After 5 days of culture, the percentage of regulatory T cells (CD3^+^CD4^+^CD25^+^FOXP3^+^) was determined by flow cytometer (Beckmann Coulter, US) and analyzed using Kaluza Software Version 2.1 [18].

### 10. Measurement of intracellular Reactive oxygen species (ROS)

Intracellular ROS level was measured in cells using MitoSOX red according to the manufacturer’s instructions. Briefly, HUH7 cells were seeded in a 6-well plate at a confluence of 1×10^6^ cells/well. Cells were exposed to 100 μM H_2_O_2_ for 30 minutes. Then cells were washed twice with 1X PBS and treated with 50 µg/mL V-BM-sEVs, V-WJ-sEVs, Apo-BM-sEVs, and Apo-WJ-sEVs for 48 hours. After treatment, cells were incubated with MitoSOX red at a concentration of 4 μM for 20 minutes in respective media at 37 °C with 5% CO2 in the dark. The quantification of ROS in terms of the percentage of positive cells was done using a flow cytometer and analyzed using Kaluza Software Version 2.1.

### 11. Cellular Bioenergetics studies

Oxygen consumption rate (OCR) was measured using Seahorse XFe24 Extracellular Flux Analyzer (Agilent Technologies, California, USA) serving as an indicator for OXPHOS. HUH7 cells were seeded at a consistent density of 1 × 10^5^/well on CellTak -coated plates (Corning) in XF-Base Media (Agilent Technologies) containing 2.5 mM glucose, 1 mM sodium pyruvate, and 2 mM glutamine and co-cultured with V-BM-sEVs, V-WJ-sEVs, Apo-BM-sEVs, and Apo-WJ-sEVs for 48h. OCR was measured sequentially at the basal level followed by the addition of 1.0 mM oligomycin, 0.75 mM FCCP (fluorocarbonyl cyanide phenylhydrazone), and 0.5 mM rotenone + antimycin to evaluate the changes in mitochondrial respiratory parameters. The plate underwent three measurements at both baseline and post-drugs addition [25]. Data were analyzed using Wave 2.6.1 and GraphPad Prism.

### 12. Quantitative real-time polymerase chain reaction (qRT-PCR)

#### miRNA expression

Total RNA was extracted from the V-BM-sEVs, V-WJ-sEVs, Apo-BM-sEVs, Apo-WJ-sEVs using Qiazol Reagent and miRNA was isolated using miRNeasy mini kit as per manufacturer’s protocol. The miRNA concentration and purity were measured by a Nanodrop 2000 Spectrophotometer (Thermo Scientific, USA). cDNA was prepared using a PCR starter kit. qRT-PCR was performed in triplicate with a PCR starter kit in CFX96 Real-Time System (Bio-Rad, USA). hsa-miR-16-5p was used as an internal control to normalize the gene expression. hsa-miR-125b-5p, hsa-miR-145b-5p, hsa-miR-34a, and miR-21 expression levels were determined using the 2-ΔΔCt method.

#### RNA expression

For the isolation of total RNA from THP1 cells after treatment with V-BM-sEVs, V-WJ-sEVs, Apo-BM-sEVs, Apo-WJ-sEVs, the total RNA was isolated using the phenol-chloroform extraction method using TRIzol reagent according to the manufacturer’s protocol. Then, 2 μg total RNA was reverse transcribed to give cDNA using the High-Capacity cDNA Reverse Transcription Kit. Quantitative real-time polymerase chain reaction (qRT-PCR) was performed using a CFX96 Real-Time System (Bio-Rad). Glyceraldehyde-3 phosphate dehydrogenase (GAPDH) was used as the housekeeping gene to normalize the gene expression. qRT-PCR was performed in triplicate using SYBR Green Master Mix according to the manufacturer’s instructions. The comparative 2−ΔΔCt method was performed to evaluate the mRNA expression of IL-1β, TNF-α and IL-10 [22].

### 13. Statistical analysis

In this study, all statistical analyses were conducted through GraphPad Prism 8.4.3 software. An unpaired student’s *t-*test was used to compare the two groups. One-way and Turkey’s post hoc tests were used to compare three or more groups. A p-value of <0.05 was considered statistically significant.

## Results

### MSCs Retain Their Characteristics Upon Apoptotic Induction

MSCs isolated from Bone marrow and Wharton’s jelly exhibited all characteristic features as per International Society of Cell and Gene Therapy (ISCT) guidelines including plastic adherence, fibroblast-like spindle morphology, trilineage differentiation, presence of surface markers CD29, CD90, CD73, CD105, HLA I, and the absence of CD34, CD45, and HLA II (Figure S1).

**Figure S1:**
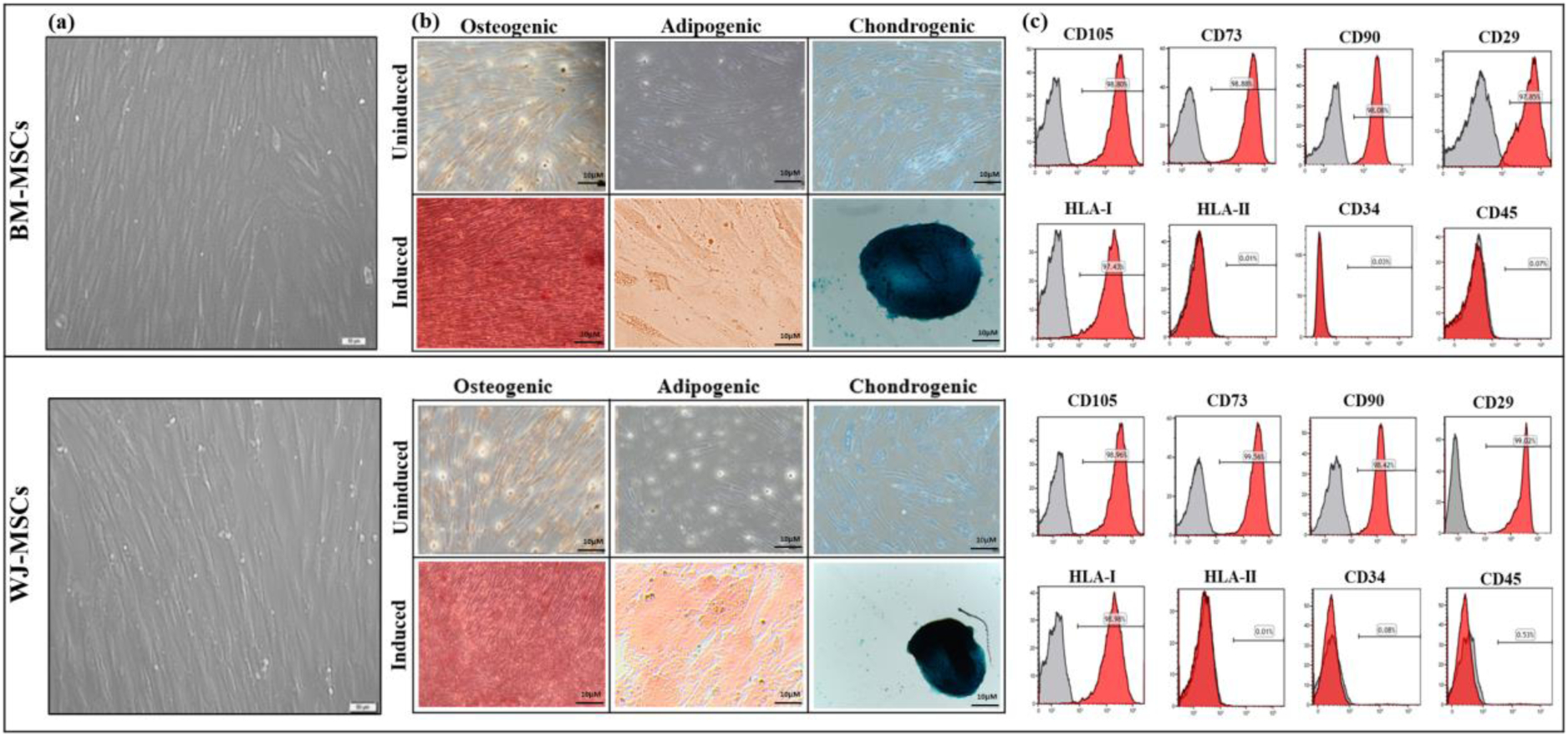
Characterization of tissue-specific (BM/WJ) human MSCs as per ISCT guidelines (a) Representative images of fibroblast-like morphology of Bone Marrow and Wharton’s Jelly MSCs (Scale bar: 100µm). (b) Trilineage differentiation of Bone Marrow and Wharton’s Jelly MSCs into Osteocytes(21 days of induction), Adipocytes (28 days of induction), and Chondrocytes (14 days of induction) (Scale bar: 100µm). (c) Surface marker profiling of Bone Marrow and Wharton’s Jelly MSCs via flow cytometry.

MSCs from both sources were subjected to apoptotic induction using 0.5 μM STS for 6h, 12h, and 24h. Annexin/PI staining was used to assess the percentage of cumulative apoptotic cells which was 99.45% (53.37% early & 46.08% late apoptotic cells) and 99.60% (52.50% early & 47.10% late apoptotic cells) at a 12h time point in BM-MSCs & WJ-MSCs respectively, whereas at 24h there was a decline in the number of apoptotic cells in both the groups (99.30% and 89.28% respectively) (Figure 1B, C). Furthermore, at this time point, the cells displayed the hallmark features of apoptosis induction including significant morphological alteration, cell blebbing, and shrinkage [26] (Figure 1A). Induction of apoptosis was also validated via western blotting in both the tissue-specific MSCs in terms of the Caspase 3 expression (Figure S2).

**Figure 1:**
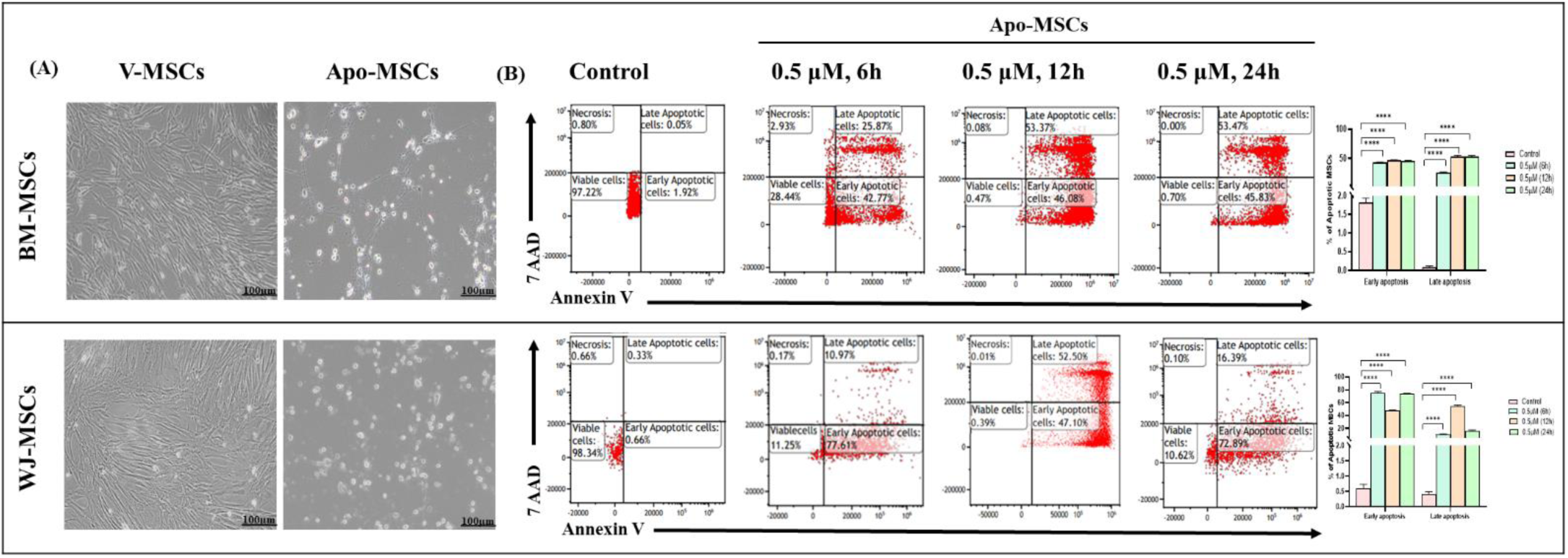
Generation of apoptotic MSCs using Staurosporine (A) Representative morphology of Apoptotic Bone Marrow and Wharton’s Jelly MSCs (Scale bar: 100µm). (B) Dot plot representing the percentage of viable, early apoptosis, late apoptosis, and necrotic cells with and without treatment of STS at a dose of 0.5 μM for 6, 12, 24 hours. (C) Bar graph depicts the percentage of apoptotic MSCs with and without treatment of staurosporine. Data shown as mean ± SD; Statistical analysis: ****p < 0.0001. All experiments were done in triplicates. Abbreviation-V: Viable; Apo: Apoptotic

**Figure S2:**
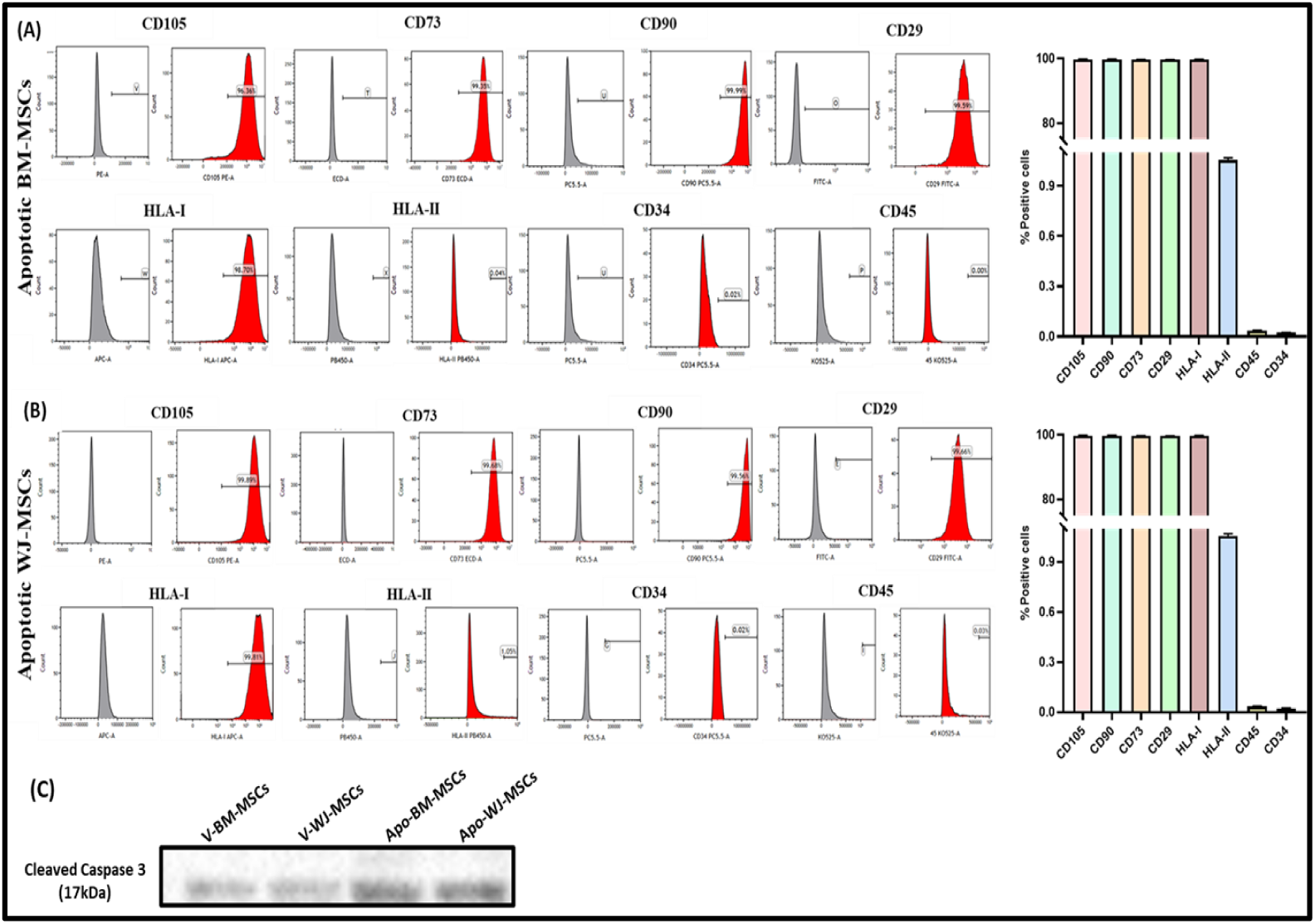
Characterization of tissue-specific Apo (BM/WJ) human MSCs as per ISCT guidelines. Flow cytometric analysis of (A) Apo-BM-MSCs and (B) Apo-WJ-MSCs for positive surface markers CD90, CD 105, CD29, CD73 and HLA-I and negative surface markers HLA-II and CD34/45. (C) Western blot shows the presence of Cleaved Caspase 3. Abbreviation-V: Viable; Apo: Apoptotic

Moreover, the immunophenotyping assay with both BM & WJ-Apo-MSCs validated the maintenance of MSCs characteristics after apoptosis induction (Figure S2).

### Apo-sEVs maintain similar characteristics to V-sEVs

sEVs obtained from the conditioned medium of both tissue-specific V-MSCs and Apo-MSCs displayed analogous characteristics in accordance with the MISEV 2018 guidelines [28]. These features encompassed a cup-shaped morphology, indicating preserved membrane integrity as observed through TEM; a size below 200nm, as identified by NTA; and the existence of surface tetraspanin CD63, cytoplasmic protein ALIX, along with the absence of Calnexin (sEVs negative marker), confirmed via western blotting (Figure 2). Additionally, cleaved caspase 3 was also detected in both tissue specific Apo-sEVs, but not in V-sEVs, confirming their respective apoptotic & viable origins [19] (Figure 2).

**Figure 2.**
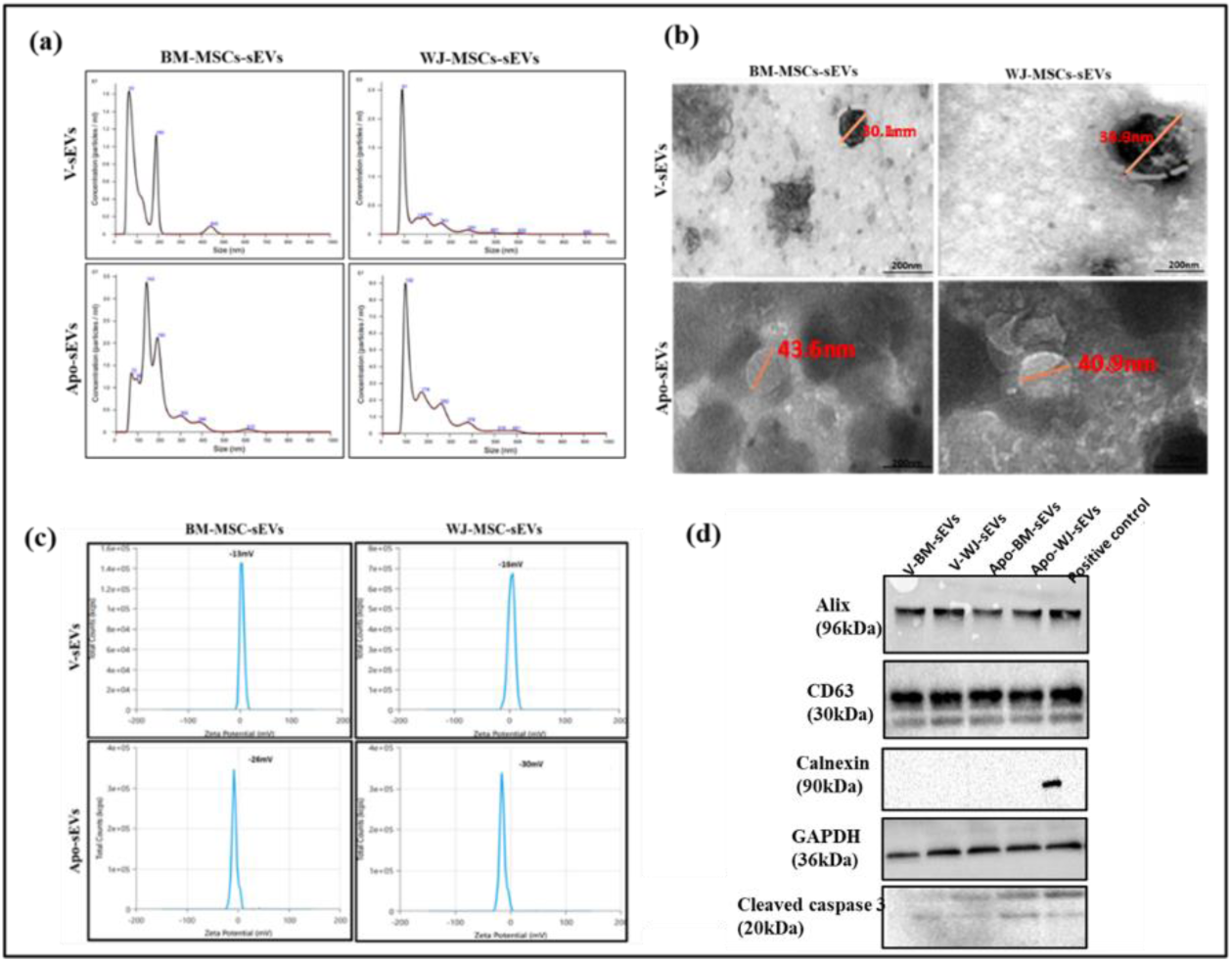
Characterization of tissue-specific (BM/WJ) human MSC-derived V-sEVs and Apo-sEVs for (a) Size distribution analysis using Nanoparticle Tracking Analysis (NTA) (b) Morphological analysis by Transmission Electron microscopy (TEM) (Scale Bar =200nm) (c) Surface charge distribution using Zeta Sizer (d) Western blot shows the presence of Alix, CD63, absence of calnexin, Cleaved Caspase 3. Abbreviation-V: Viable; Apo: Apoptotic

Dissecting into the results obtained from NTA analysis we found that Apo-BM & WJ-sEVs showed a higher particle concentration (3.5 ×10^7^ and 2.5 ×10^8^ respectively) as compared to the V-BM & WJ-sEVs (1.6 ×10^7^ and 9 ×10^7^respectively) (Figure 2B); and demonstrated a significantly greater (p: ≤0.0001, p: ≤0.01) mean particle size (143 nm and 102 nm, respectively) as compared to the V-BM & WJ-sEVs (63 nm and 91 nm, respectively).

Also, the membrane potential of Apo-BM & WJ-sEVs was −13mV and −16 mV respectively, as compared to the membrane potential of their viable counterparts −26mV and −30mV (Figure 2A, 2C, S3) as observed via the ZetaPotential Analysis.

**Figure S3.**
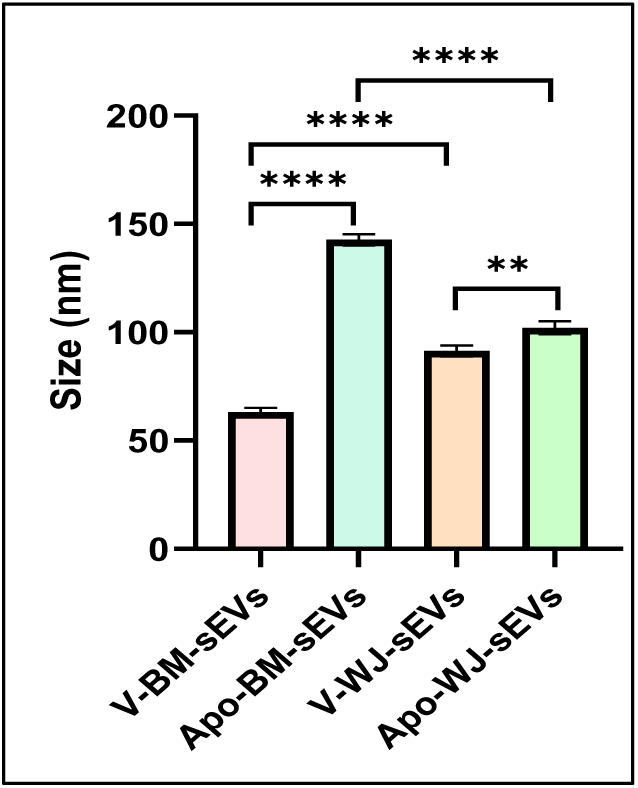
Characterization of tissue-specific (BM/WJ) human MSC-derived V-sEVs and Apo-sEVs for Size distribution analysis using Nanoparticle Tracking Analysis (NTA)

### Apo-sEVs are highly instrumental in driving macrophage polarization towards anti-inflammatory phenotype via efferocytosis

To validate their immunomodulatory functionality, LPS-activated THP1-induced macrophages were co-cultured with tissue specific V-& Apo-sEVs. Herein, we observed that there was a significant polarization of M1 macrophages towards the M2 phenotype after Apo-sEVs treatment as compared to the V-sEVs treatment in both tissue sources. The polarization was assessed in terms of a significant increase in Arginase & CD206 expression (M2 markers; p: ≤0.0001), and the concurrent decrease in iNOS (M1 marker; p: ≤0.0001) (Figure 3, S4).

**Figure 3.**
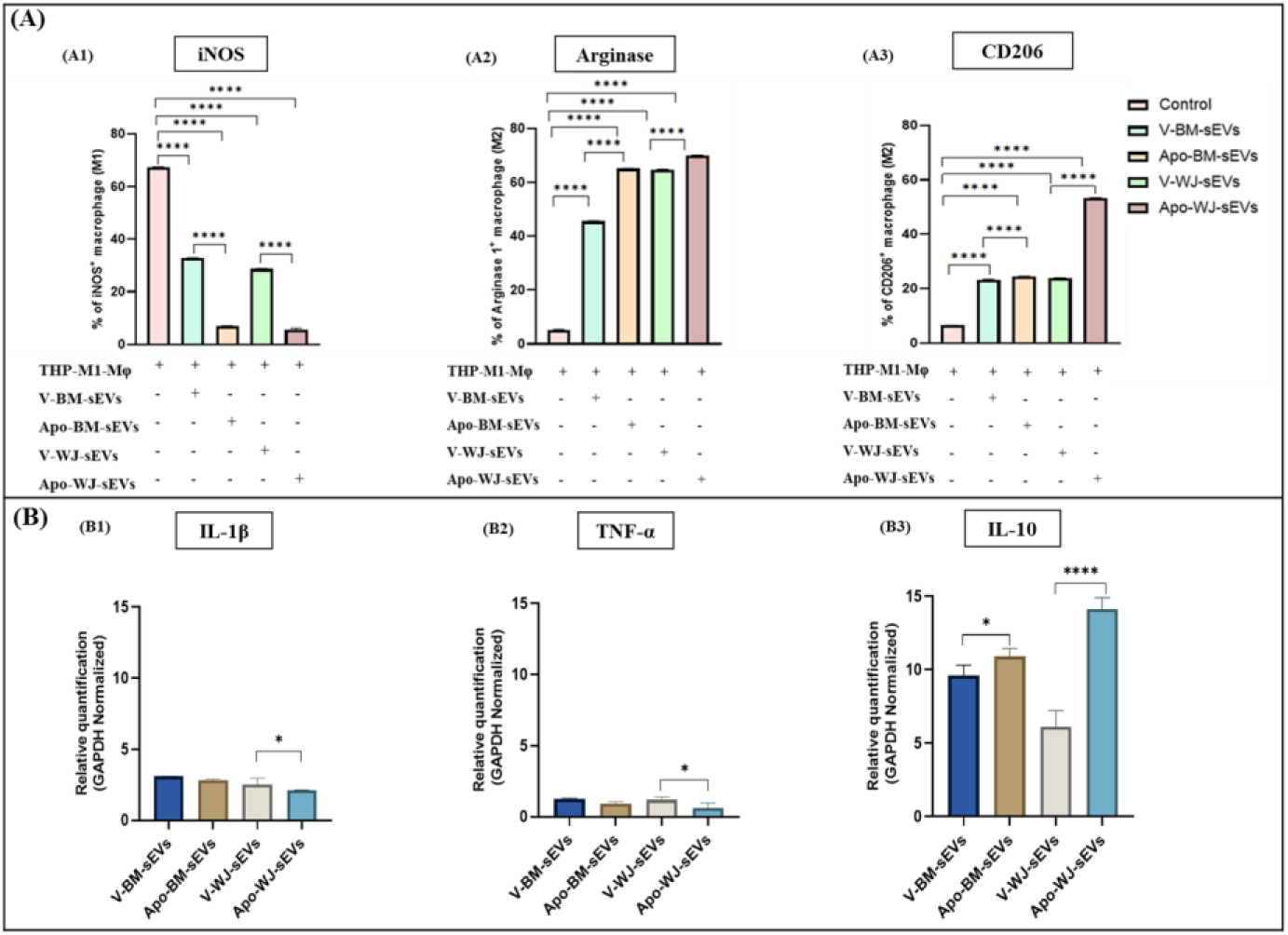
Apo-sEVs enhanced the polarization of M1 macrophages to M2 macrophages. (A) Bar graph showing the expression of (a) iNOS (M1), (b) Arginase, and (c) CD206 (M2) after the treatment of tissue-specific (BM/WJ) human MSC-derived V-sEVs and Apo-sEVs. (B) Relative mRNA expression of (a) IL-Iβ, (b) TNF-α, and (c) IL-10 after the treatment of tissue-specific (BM/WJ) human MSC-derived V-sEVs and Apo-sEVs by qP^1^C^4^R. Data shown as mean ± SD; Statistical analysis: ****p < 0.0001, **p < 0.01, *p < 0.05. All experiments were done in triplicates. Abbreviation-V: Viable; Apo: Apoptotic

**Figure S4.**
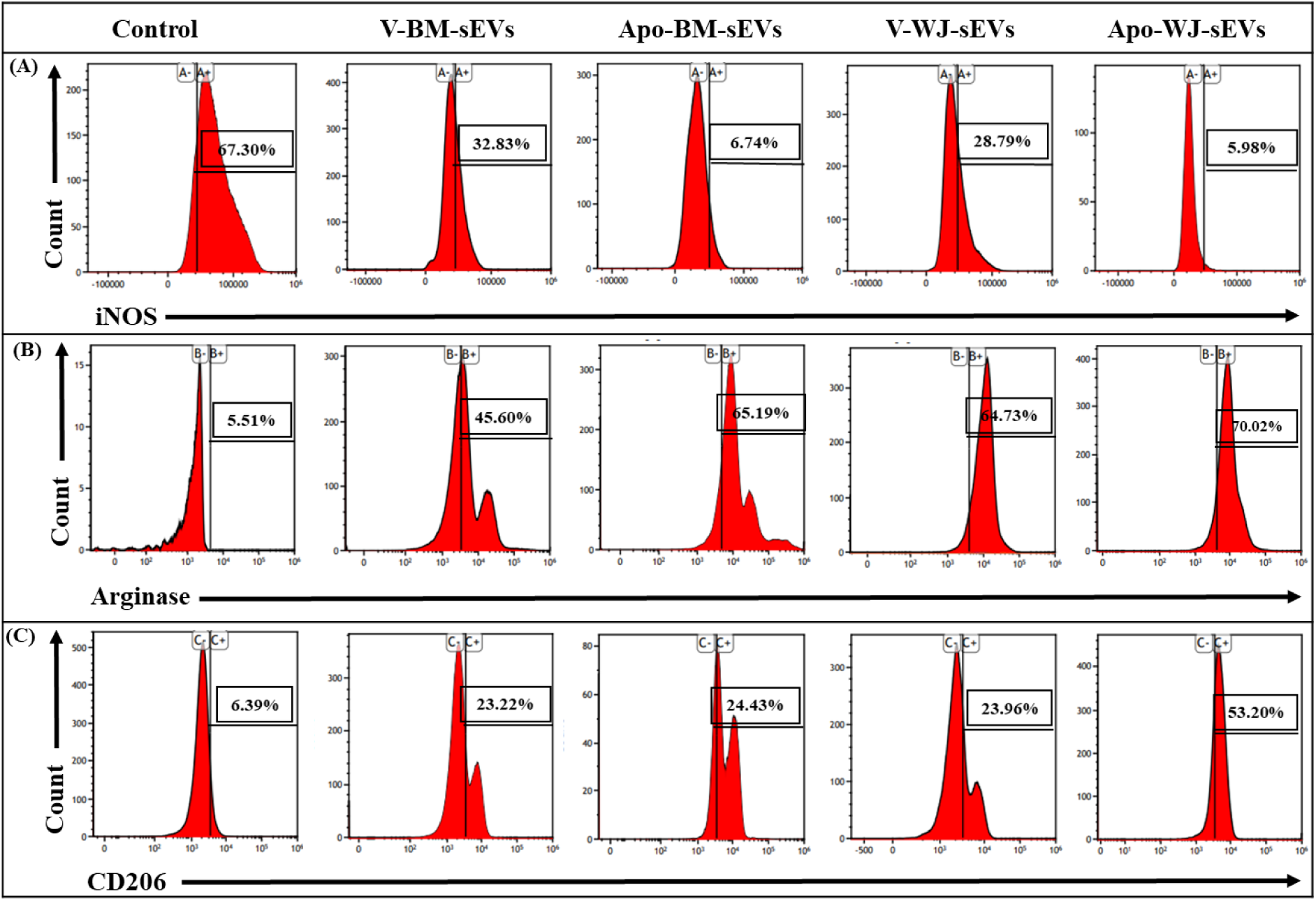
Apo-sEVs enhanced the polarization of M1 macrophages to M2 macrophages. (A) Flow cytometry analysis shows the expression of (A) iNOS (M1), (B) CD206, and (C) Arginase-1 (M2) after the treatment of tissue-specific (BM/WJ) human MSC-derived V-sEVs and Apo-sEVs.

In terms of the tissue source, it was observed that sEVs derived from Apo-WJ-MSCs were more efficient in macrophage polarization as compared to the Apo-BM-MSCs. This was ascertained by the significant increase in anti-inflammatory M2 markers Arginase (59.68%; p:≤0.0001 and 64.51%; p:≤0.0001 in Apo-BM-& Apo-WJ-sEVs respectively) & CD206 expression (24.43%; p:≤0.0001 and 53.20%; p:≤0.0001 in Apo-BM-& Apo-WJ-sEVs respectively) (Figure 3A, S4).

A similar trend was also observed in the gene expression level of proinflammatory (IL-1β & TNFα) & anti-inflammatory (IL-10) cytokines, wherein the latter was significantly decreased, and the later was significantly increased in Apo-sEVs as compared to V-sEVs (p: ≤0.001) (Figure 3B).

To understand the aforementioned heightened functional capabilities of Apo-sEVs, particularly those derived from Apo-WJ-MSCs, we assessed the efferocytotic activity of both tissue-specific V- and Apo-sEVs. The results revealed that Apo-BM & WJ-sEVs exhibited significantly increased efferocytosis activity (73.13% and 94.93%; p: ≤0.001 respectively) compared to V-BM & WJ-sEVs (29.06% and 36.68%, respectively), thereby explaining the higher immunomodulation via Apo-WJ-sEVs (Figure 4).

**Figure 4:**
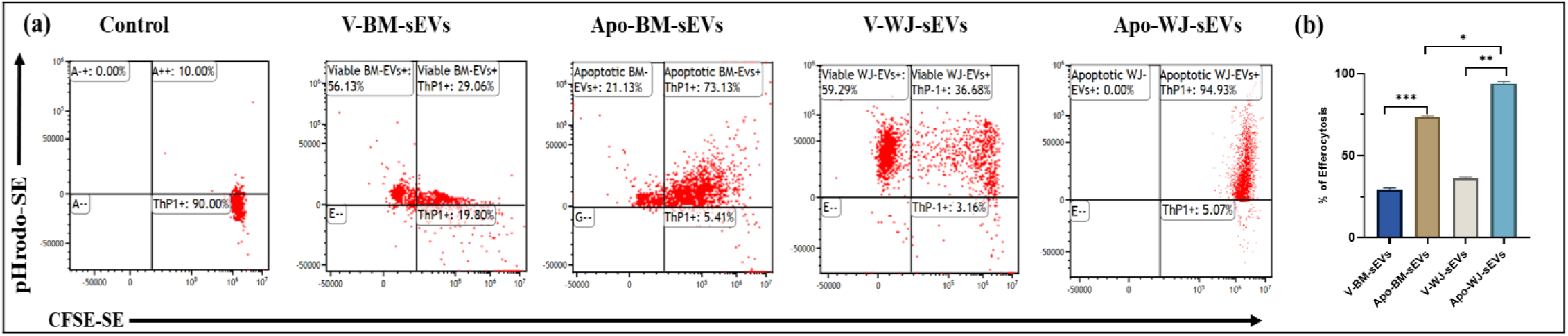
Efferocytosis of Apo-sEVs enhanced by macrophages. A representative (a) dot plot and (b) bar graph showing the magnitude of efferocytosis of tissue-specific (BM/WJ) human MSC-derived V-sEVs and Apo-sEVs by macrophage. Data shown as mean ± SD; Statistical analysis: ***p < 0.001, **p < 0.01, *p < 0.05. All experiments were done in triplicates. Abbreviation-V: Viable; Apo: Apoptotic

### Apo-sEVs Suppress T-Cell Proliferation and Enhance Treg Induction as one of their Immunomodulatory Mechanisms

For the evaluation of the immunomodulatory potential of Apo-sEVs, we investigated their role on the proliferation of immune-inductive T cells. Additionally, we assessed whether Apo-sEVs could induce a more effective polarization of T cells towards regulatory T cells (Tregs) compared to V-sEVs [18].

It was found that Apo-sEVs significantly reduced CD3^+^ T cell proliferation & enhanced their induction into Tregs as compared to V-sEVs in both the tissue sources (p: ≤0.0001). Furthermore, Apo-WJ-sEVs were able to confer a greater reduction in T-cell proliferation as compared to the Apo-BM-sEVs (63% & 68.59%; p: ≤0.0001 respectively) (Figure 5A1, A2). Moreover, Treg induction was significantly enhanced in Apo-WJ-sEVs (22.12%) as compared to Apo-BM-sEVs (17.10%) (Figure 5B1, B2).

**Figure 5:**
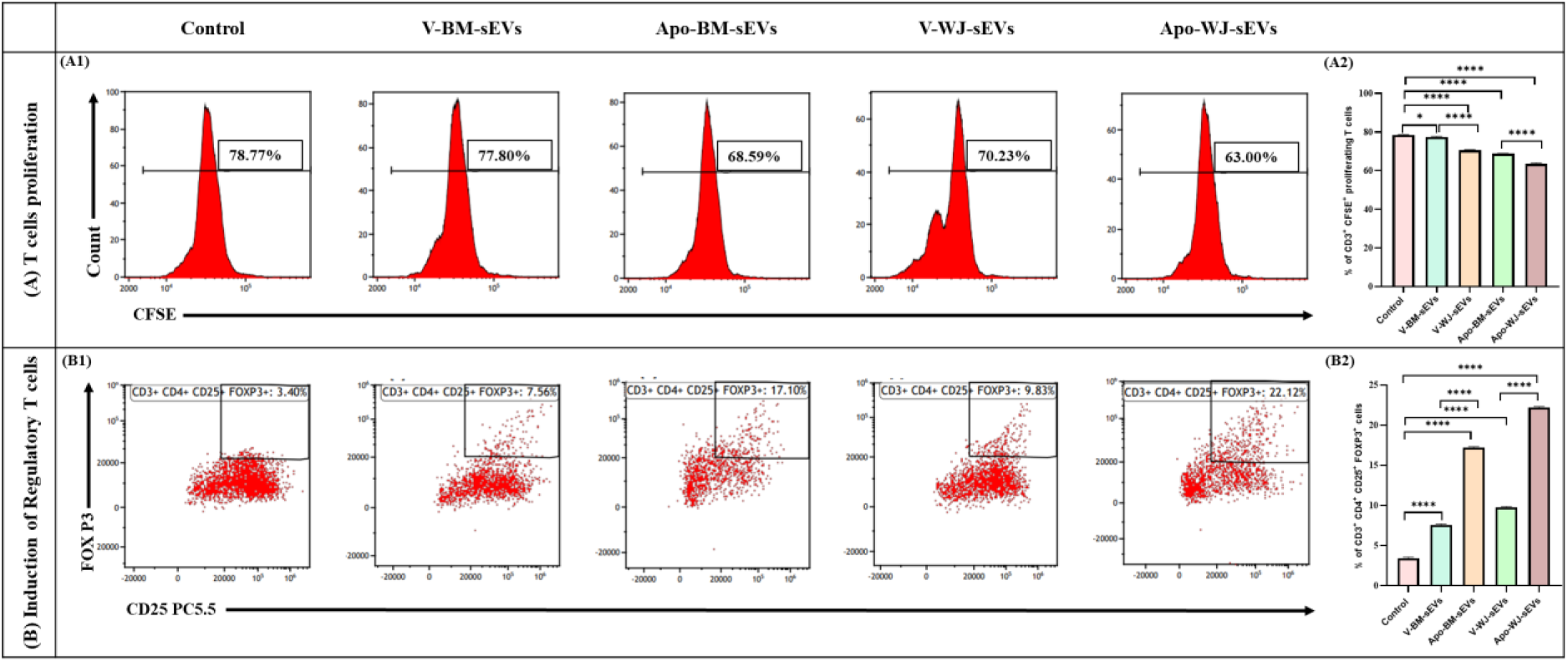
Apoptotic sEVs mediated suppression of T cell proliferation and induction of regulatory T cells. (A) Immunosuppressive effect of tissue-specific (BM/WJ) human MSC-derived V-sEVs and Apo-sEVs on the proliferation of T cells using CFSE-based T cell proliferation assay. (B) Induction of FoxP3 Regulatory T cells after treatment with tissue-specific (BM/WJ) human MSC-derived V-sEVs and Apo-sEVs. Data are shown as mean ± SD; Statistical analysis: ****p < 0.0001. All experiments were done in triplicates. Abbreviation-V: Viable; Apo: Apoptotic

### Efficient Mitochondrial ROS alleviation by Apo-sEVs as compared to their Viable Counterparts

For the evaluation of oxidative stress reduction potential of sEVs, we treated the hydrogen peroxide-induced HUH7 cells with the tissue specific V-sEVs & Apo-sEVs for 24h, post which the oxidative stress was evaluated using Mitosox staining via flow cytometry. Our results indicated that oxidative stress was significantly reduced in Apo-BM-sEVs (38%, p: ≤0.01) and Apo-WJ-sEVs (34%, p: ≤0.001) treated groups as compared with the V-BM-sEVs (45%, p: ≤0.05%) and V-WJ-sEVs (42%, p: ≤0.001) treated groups (Figure 6, S5). This demonstrated that apoptotic induction to the MSCs can significantly enhance the reparative capabilities of sEVs derived from them, and inhibit the oxidative stress induced injury [45,46].

**Figure 6.**
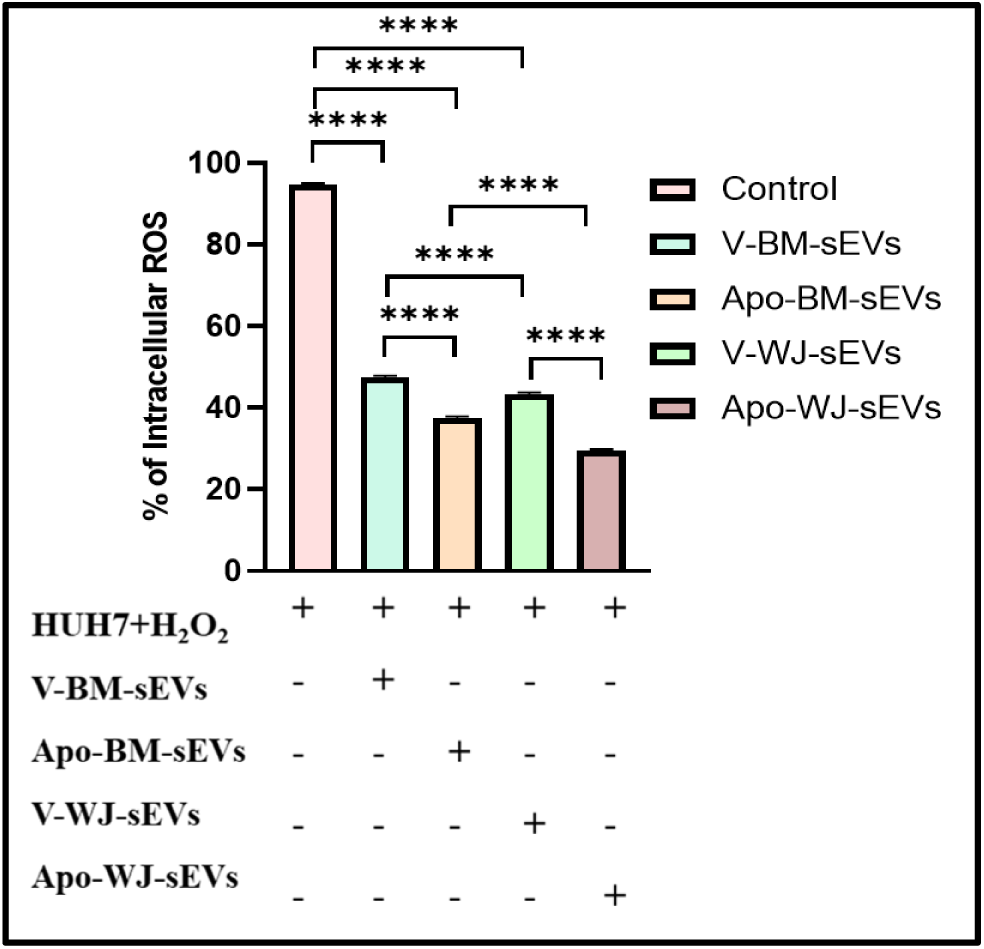
Regulation of Intracellular ROS level in HUH7 cells after Apo-sEVs treatment. Quantitation of intracellular ROS levels in HUH7 cells after the treatment of tissue-specific (BM/WJ) human MSC-derived V-sEVs and Apo-sEVs by qPCR. Data shown as mean ± SD; Statistical analysis: ****p < 0.0001. All experiments were done in triplicates. Abbreviation-V: Viable; Apo: Apoptotic

**Figure S5.**
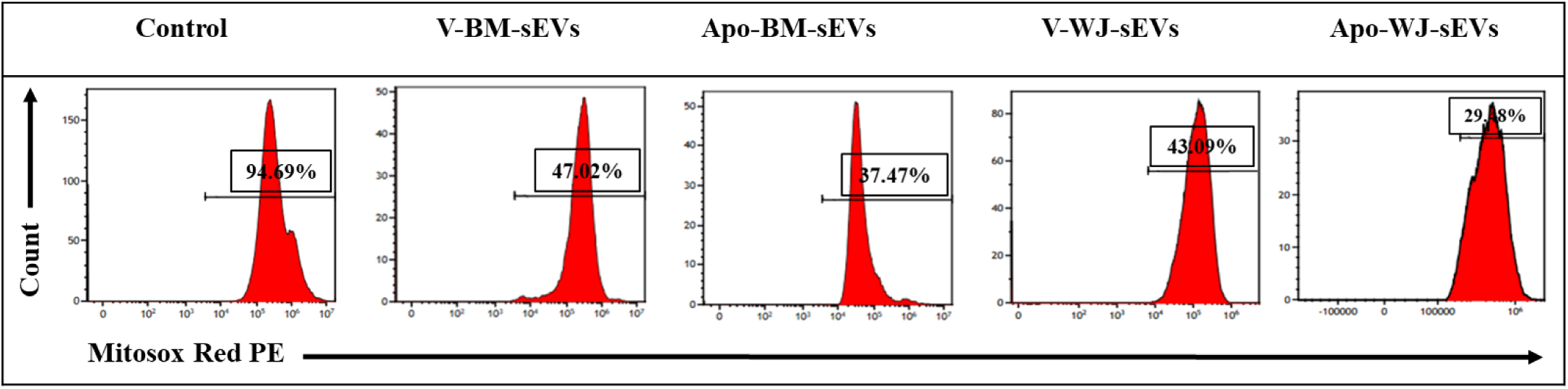
Regulation of Intracellular ROS level in HUH7 cells after Apo-sEVs treatment. Flow cytometry analysis shows the reduction of intracellular ROS levels in HUH7 cells after the treatment of tissue-specific (BM/WJ) human MSC-derived V-sEVs and Apo-sEVs.

In order to further specifically analyse the effect of both V-sEVs & Apo-sEVs on mitochondrial health, we performed the seahorse extracellular flux mitochondrial stress assay. Upon treatment of HUH7 cells with sEVs from all groups (tissue specific; viable and apoptotic MSCs), it was observed that the basal respiration rate, the maximal respiration rate & the spare respiratory reserve were improved in contrast to the untreated group. However, concerning the mitochondrial functionality, as assessed by the ATP production capacity and the proton leak, it was observed that Apo-sEVs were significantly improving the ATP production & reducing the proton leak in the cells as compared to the V-sEVs, regardless of the tissue source. It was also found that Apo-WJ-sEVs were observed to significantly enhance basal respiration as compared to other groups (p: ≤0.05) (Figure 7), as compared to Apo-BM-sEVs.

**Figure 7.**
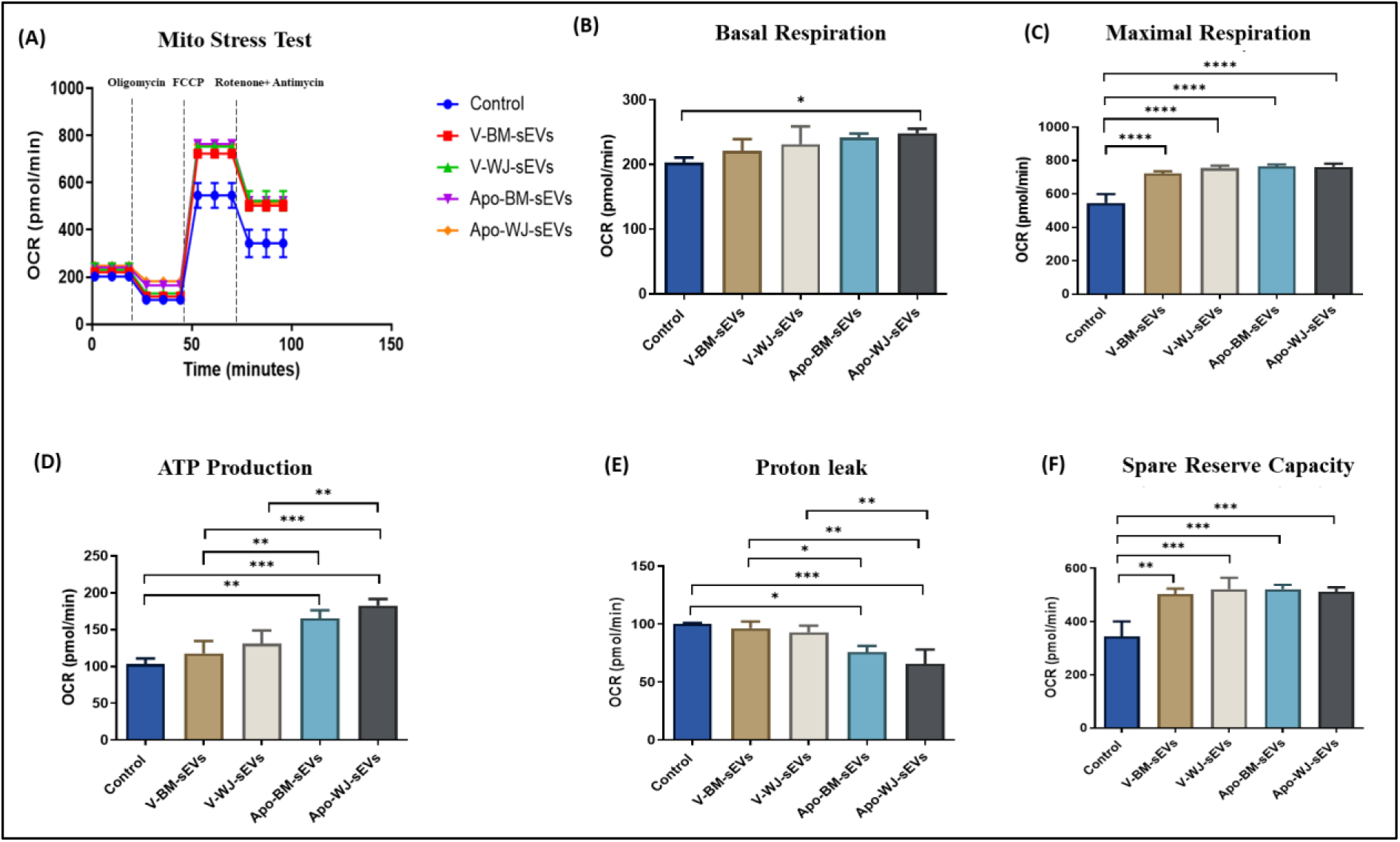
Mitochondrial parameters related to bioenergetics of tissue-specific Apo-MSC-sEVs. (A) Real-time changes in oxygen consumption rate (OCR) with subsequent treatment with oligomycin, FCCP and rotenone and antimycin A. Apo-sEVs improves the mitochondrial respiratory parameters including (B) Basal respiration, (C) Maximal respiration (D) ATP production, (E) Proton Leak, (F) Spare Reserve capacity of HUH7 cells. Data shown as mean ± SD; Statistical analysis: ***p < 0.001, **p < 0.01, *p < 0.05. All experiments were done in triplicates. Abbreviation-V: Viable; Apo: Apoptotic

### Immunomodulatory miRNAs are Enriched in Apo-sEVs

To identify the mechanism for the enhanced immunomodulatory & antioxidant functionality of Apo-sEVs, we sought to evaluate the content of Apo-sEVs in comparison to V-sEVs, in terms of miRNAs known to target antioxidant & immunomodulatory signaling pathways.

Therefore, we assessed the expression of miR125b-5p, miR145a-5p, miR21-5p, and miR34 in both the tissue-specific V-sEVs and Apo-sEVs via qPCR. The results showed a significant increase in the packaging of all 4 regenerative miRNAs in Apo-sEVs in comparison with V-sEVs in both BM & WJ (p: ≤0.001). Specifically, there was a 3.5-, 3.2-, 9-, and 5-fold higher upregulation of miRNA 125b-5p, miR 145b-5p, miR 21, and miR 34 respectively in Apo-WJ-sEVs as compared to Apo-BM-sEVs (Figure 8).

**Figure 8.**
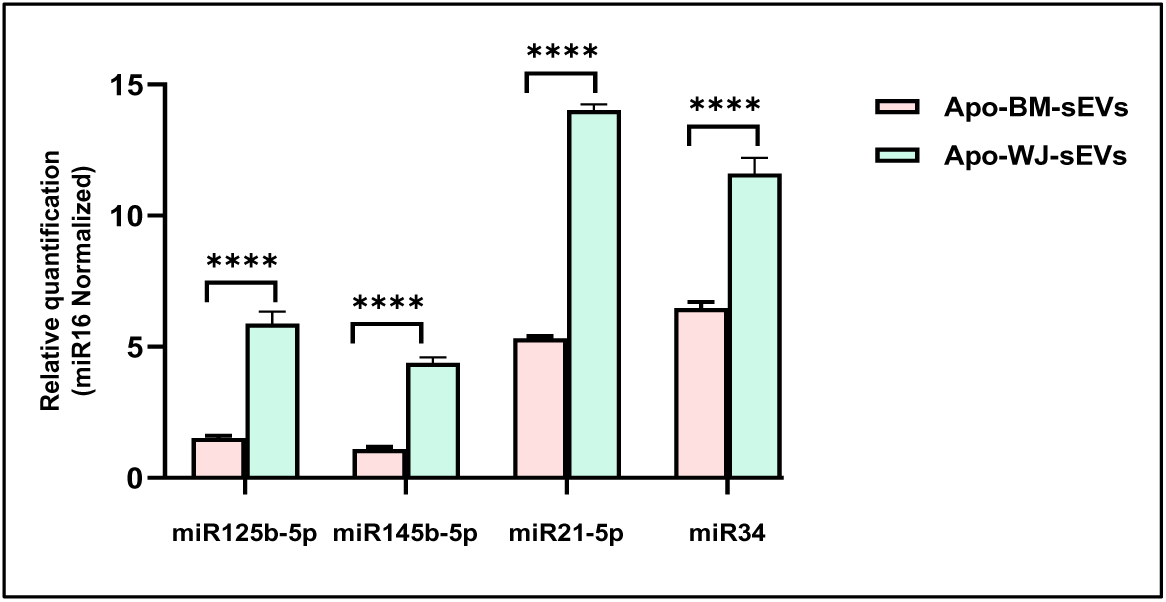
Upregulated expression of hepatoprotective miRNA in Apo-sEVs. Quantitative analysis of miR 125b-5p, miR 145b-5p, miR 21-5p, miR 34a in tissue-specific (BM/WJ) human MSC-derived V-sEVs and Apo-sEVs by qPCR. Data shown as mean ± SD; Statistical analysis: ****p < 0.0001. All experiments were done in triplicates. Abbreviation-V: Viable; Apo: Apoptotic

## Discussion

Mesenchymal stem cells (MSCs) are currently leading the way in regenerative medicine therapeutics [66]. Within their natural niche, MSCs demonstrate regenerative capabilities through paracrine secretions [67]. Moreover, several reports indicate that MSCs undergo apoptosis post-infusion to exhibit their reparative functions [11]. Consequently, few studies have investigated the role of Apo-EVs in regeneration [22]. In this context, our study assessed the immunomodulatory and ROS alleviation potential of tissue-specific Apo-MSC-sEVs, comparing them with V-sEVs, using an in vitro liver injury model.

Both BM & WJ-MSCs were induced to undergo apoptosis using 0.5 μM STS, which has also been previously tested by Juan Wang et al, 2021 in their study to induce apoptosis in MSCs [19,20]. However, due to variability in cellular responses, we evaluated apoptosis induction in BM & WJ-MSCs at different time points, and analyzed the extent of apoptosis induction. Our observations showed that both cell types were most receptive to this induction strategy at 12 hours, where they also displayed morphological alterations as a result of the active cleavage of Caspase 3, as reported by other studies [11, 27].

Following apoptotic induction, we collected conditioned media to specifically isolate a homogenous population of small EVs (sEVs) with a size less than 200nm. Notably, existing literature has often encompassed the Apo-EVs population, including both small and large EVs, as well as apoptotic bodies [22]. To our knowledge, our study is the first to focus specifically on the Apo-sEVs population.

These Apo-sEVs were found to maintain the characteristics as per the MISEV guidelines; an interesting finding during the NTA analysis was however, the increased average particle size and total concentration of sEVs derived from Apo-MSCs & V-MSCs in both tissue sources, considering an identical number of MSCs (1×10^5^) and volume of conditioned media (3 ml) to start with. We speculate that the increase in size could be due to the enrichment in cargo thereby conferring a heightened functionality to Apo-sEVs over V-sEVs, as observed by subsequent *in vitro* assays as well. This is further in line with the previous finding by Liang et al. (2017), who confirmed the increase in size of HEK 293T-sEVs upon miR26a enrichment by transfection [29]. As per our current understanding, this is the first study reporting the increase in size of Apo-sEVs possibly due to cargo enrichment.

Furthermore, the increase in particle number is consistent with the previous research indicating that apoptotic cells release a greater number of microparticles compared to viable cells [18]. The speculated mechanism for the increased secretion has been suggested to involve the activation of scramblases during apoptosis, triggering membrane budding. Caspase-3 has been found to cleave Rho-associated protein kinase (ROCK I), which leads to the remodelling of actin and microtubules present at the plasma membrane, therefore resulting in membrane blebbing and EV formation. However, the precise mechanism for the heightened secretion remains unclear [26,30].

Since MSCs are widely recognised for their immunomodulatory properties, we sought to evaluate the efficacy of these sEVs in polarization of macrophages, efferocytotic activity, T-Cell Proliferation, as well T-Regulatory cells induction [18,32–34,37–38].

Macrophages play a pivotal role in initiating and resolving inflammatory responses [18]. They can adopt a pro-inflammatory M1 phenotype or shift towards an anti-inflammatory M2 phenotype in response to specific signals [27–30]. A well-established mechanism through which MSCs-sEVs exert immunomodulation is by influencing macrophage polarization [18]. Recent evidence has suggested that MSCs & their EVs undergo efferocytosis via engulfment by macrophages, thereby instructing their polarization [32–34]. These EVs are acknowledged for transporting a multitude of miRNAs that contribute to directing macrophage polarization towards the M2 phenotype. Several studies have shown that miRNAs, including miR125, miR145, miR146, miR-21, and miR223, can regulate macrophage polarization by suppressing various inflammatory targets such as Notch1, IRAK1, TRAF6, IRF5, C/EBPβ, pknox. This ultimately inhibits the NF-κB and TLR-4 driven release of proinflammatory cytokines in macrophages [35].

Our study demonstrated that Apo-sEVs were more efficient in polarization of macrophages, as well as efferocytotic activity, as compared to their viable counterparts. Moreover, WJ as a source was faring significantly better than BM. Patil et al., (2021) also observed the same phenomenon in myocardial ischemic injury, wherein they observed that MSCs-EVs were able to enhance the efferocytosis of apoptotic cardiomyocytes, thereby leading to an increased macrophage polarization activity towards the anti-inflammatory phenotype [18,36].

Prior research has identified the involvement of MSCs-sEVs in inhibiting T-cell proliferation (CD3^+^) and promoting their differentiation into regulatory T cells (CD3^+^ CD4^+^ CD25^+^ FoxP3^+^) [37,38]. This was also found to be replicated as a mechanism of action via Apo-sEVs, wherein WJ as a source was found to be more potent. This was also in line with a previous finding by Chen et al., 2019, who observed in their study that the administration of Apo-EVs derived from primary murine thymocytes & Jurkat cells were able to promote regulatory T cell induction & suppress the production of T-helper cells [18]

Both of the above compelling findings pertaining to the potential of Apo-sEVs towards macrophage polarization & T-reg induction, affirm the superior immunomodulatory potential of Apo-sEVs when compared to their viable counterparts. This robust outcome not only underscores the pivotal role of Apo-sEVs in immunomodulation over V-sEVs, but also emphasizes the heightened tissue-specific functionality of WJ-MSCs over BM-MSCs as a source.

One of the major pathophysiological features of liver diseases is the induction of reactive oxygen species (ROS) which results in oxidative stress in the cell [68]. ROS induction disrupts the electron transport chain in mitochondria, leading to disbalance in cellular homeostasis [39,40]. Multiple studies have reported the antioxidant effect of MSCs-sEVs in alleviating oxidative damage in various diseases such as liver fibrosis, cardiovascular diseases, neurodegeneration, and renal disorder [22,41,42]. A large body of evidence has shown the capabilities of V-sEVs in ROS alleviation, however there is still a dearth of knowledge about the same with Apo-sEVs [42,45]. Therefore, we decided to evaluate the ROS alleviation capability of Apo-sEVs as compared to V-sEVs in a tissue specific manner.

We tested the same with hydrogen Peroxide induced HUH7 cells as an *in vitro* model of oxidative stress. Hydrogen peroxide is known to disrupt the cellular redox system and cause DNA damage, thereby leading to changes in the fundamental cellular function, restricting the proliferation and initiating apoptosis [43,44]. Herein we observed that Apo-WJ-sEVs exhibited notable efficacy in reducing reactive oxygen species (ROS), as assessed through mitosox staining, a technique specifically targeting mitochondrial ROS. The results of this assay indicated a significant reduction in mitochondrial ROS levels upon treatment with Apo-WJ-sEVs.

Consequently, we aimed to delve deeper into the effects of Apo-sEVs using the seahorse extracellular flux mitochondrial stress assay. This assay evaluates the key parameters of mitochondrial function by directly measuring the oxygen consumption rate (OCR) of cells in real-time and providing insights into the basal & maximal respiration rate, ATP Production, Proton Leak & Spare respiratory Reserve. Multiple studies have confirmed that MSCs and MSC-EVs are instrumental in modulating the mitochondrial bioenergetics of the cells, which also leads to improved mitochondrial efficiency & regeneration [47–51].

Through this assay, we found that all the sEVs were equally efficient in improving the mitochondrial fitness, however Apo-sEVs were more efficient in reducing the proton leak & enhancing the ATP production, regardless of the tissue source. Thereby suggesting that Apo-sEVs are more potent in enhancing mitochondrial energetics, functionality, and reducing mitochondrial damage. An interesting finding further supported the supremacy of Apo-WJ-sEVs in regeneration, as they were able to significantly enhance the basal respiration rate, thereby suggesting that Apo-WJ-sEVs enable the recipient cells to meet their energy needs for carrying out reparative activity [44,45].

To this point, our investigation established the superior cumulative antioxidant and immunomodulatory capabilities of Apo-WJ-sEVs. Moreover, earlier observations revealed a notable increase in the size of Apo-sEVs, a phenomenon we hypothesized to be linked to miRNA cargo enrichment, as suggested by Liang G et al, 2018 [25]. Consequently, we analyzed the miRNA content within Apo-sEVs compared to V-sEVs in a tissue-specific manner, aiming to elucidate potential mechanisms contributing to enhanced functionality. This analysis focused on the encapsulation of specific miRNAs known to play crucial roles in immunomodulation and antioxidant activities. Previous reports have shown the packaging of miRNA 125b-5p, miRNA 145b-5p, miRNA 21-5p, and miRNA 34 in tissue-specific V-sEVs [2,40,53–55] which have established role in immunomodulation and ROS alleviation by targeting cellular signaling pathways including NFkB, JAK/STAT, MAPK & NOTCH. miR125b targets MAPK pathway-related proteins (p38, ERK1/2, and JNK1/2) [45]; miR145 targets NF-κB related proteins (OPG and KLF5) [46]; miR21 targets JAK-STAT pathway related protein (STAT3) [47]; and miR34 targets NOTCH receptor (DllA, Jag1b*, and* Jag2) [48]; thereby mitigating inflammation and oxidative stress respectively. We found that the enrichment of all these miRNAs was significantly upregulated in Apo-WJ-sEVs.

Our current findings align with this knowledge, providing further evidence that Apo-WJ-sEVs, which exhibit a higher abundance of these miRNAs, possess an enhanced potential for immunomodulation & ROS alleviation [47–50] It demonstrates the potential of apoptosis induction as a strategy to enhance the immunomodulatory & regenerative capabilities of MSCs derived sEVs. It also presents an approach towards developing an economical and minimally manipulated regenerative modality. Although the current findings were validated in an *in vitro* liver injury model, its impact is far and wide, showing hope for other degenerative diseases. While existing literature primarily focuses on assessing the regenerative and immunomodulatory abilities of V-sEVs, our study stands out as the first to compare the regenerative potential of Apo-sEVs across 2 different MSC tissue sources-Bone Marrow & Wharton’s Jelly. Notably, Apo-WJ-sEVs faired significantly better in all the domains of *in vitro* functionality evaluated in the current study. Furthermore, our study is the first to focus on a homogeneous population of sEVs (size <200nm), eliminating the vesicles possibly having detrimental components as a by-product of apoptosis. Our findings highlight a manifold enhancement in the therapeutic and regenerative potential of Apo-sEVs, positioning them as a promising cell-free therapy, especially in conditions where MSCs penetration is challenging, such as brain and heart disorders. The increased yield of sEVs upon apoptotic induction further emphasizes its potential for translation, addressing the significant roadblock of low sEVs yield in clinical applications. However, further investigations into the specific mechanisms, contents, and *in vivo* functionality of Apo-sEVs in various disease scenarios are crucial to unlocking their clinical potential. Nonetheless, this study acts as a stepping stone for further investigation in this domain, underscoring the necessity for exploration and research to delve into the underlying mechanisms behind the regenerative and immunomodulatory potential of Apo-sEVs.

## Acknowledgments

The authors acknowledge the Sophisticated Analytical Instrumentation Facility, AIIMS, New Delhi, India for their technical support in Electron Microscopy imaging. We also sincerely acknowledge Dr. Amit Kumar Dinda for providing the NTA Facility. The graphical abstract was created using Biorender.com.

## Conflict of Interest

The authors declare no conflict of interest.

## Author Contributions

SM was involved in the study conception, designing and supervision of experiments, interpretation of data, and drafting the manuscript. MM and MM were involved in performing the experimental procedures, data acquisition, analysis, and interpretation, and writing the manuscript. YS contributed to the interpretation of data and drafting of the manuscript. RKS and NM are involved in the interpretation of data. All authors approved the final version of the manuscript.

## Ethics Statement

The study was commenced after obtaining ethical approval from the Institutional Ethical Committee (IEC) and Institutional Committee for Stem Cell Research (IC-SCR), AIIMS, New Delhi, India.

## Funding statement

The authors express their gratitude to the All-India Institute of Medical Sciences (AIIMS), New Delhi, Department of Biotechnology, Government of India, New Delhi (Grant No.BT/01/COE/07/03) for their assistance in supporting this manuscript. MM extends acknowledgment to the Department of Science & Technology (DST), Ministry of Science and Technology, Government of India, for providing the fellowship.

## Notes

### Competing Interest Statement

The authors have declared no competing interest.

## References

1. Rawat S, Srivastava P, Prabha P, Gupta S, Kanga U, Mohanty S. A comparative study on immunomodulatory potential of tissue-specific hMSCs: Role of HLA-G. 2018 Jun 1;

2. Soni N, Gupta S, Rawat S, Krishnakumar V, Mohanty S, Banerjee A. MicroRNA-Enriched Exosomes from Different Sources of Mesenchymal Stem Cells Can Differentially Modulate Functions of Immune Cells and Neurogenesis. Biomedicines. 2021 Dec 30;10(1):69.

3. Zanirati G, Provenzi L, Libermann LL, Bizotto SC, Ghilardi IM, Marinowic DR, et al. Stem cell-based therapy for COVID-19 and ARDS: a systematic review. Npj Regen Med. 2021 Nov 8;6(1):1–15.

4. Li Y, Hao J, Hu Z, Yang YG, Zhou Q, Sun L, et al. Current status of clinical trials assessing mesenchymal stem cell therapy for graft versus host disease: a systematic review. Stem Cell Res Ther. 2022 Mar 4;13(1):93.

5. Jurado M, De La Mata C, Ruiz-García A, López-Fernández E, Espinosa O, Remigia MJ, et al. Adipose tissue-derived mesenchymal stromal cells as part of therapy for chronic graft-versus-host disease: A phase I/II study. Cytotherapy. 2017 Aug 1;19(8):927–36.

6. Cheung TS, Giacomini C, Cereda M, Avivar-Valderas A, Capece D, Bertolino GM, et al. Apoptosis in mesenchymal stromal cells activates an immunosuppressive secretome predicting clinical response in Crohn’s disease. Mol Ther. 2023 Dec 6;31(12):3531–44.

7. Preda MB, Neculachi CA, Fenyo IM, Vacaru AM, Publik MA, Simionescu M, et al. Short lifespan of syngeneic transplanted MSC is a consequence of in vivo apoptosis and immune cell recruitment in mice. Cell Death Dis. 2021 Jun 2;12(6):1–12.

8. Takakura Y, Matsumoto A, Takahashi Y. Therapeutic Application of Small Extracellular Vesicles (sEVs): Pharmaceutical and Pharmacokinetic Challenges. Biol Pharm Bull. 2020;43(4):576–83.

9. Charoenviriyakul C, Takahashi Y, Nishikawa M, Takakura Y. Preservation of exosomes at room temperature using lyophilization. Int J Pharm. 2018 Dec 20;553(1):1–7.

10. Wu JY, Li YJ, Hu XB, Huang S, Xiang DX. Preservation of small extracellular vesicles for functional analysis and therapeutic applications: a comparative evaluation of storage conditions. Drug Deliv. 2021 Jan 1;28(1):162–70.

11. Pang SHM, D’Rozario J, Mendonca S, Bhuvan T, Payne NL, Zheng D, et al. Mesenchymal stromal cell apoptosis is required for their therapeutic function. Nat Commun. 2021 Nov 11;12(1):6495.

12. He X, Hong W, Yang J, Lei H, Lu T, He C, et al. Spontaneous apoptosis of cells in therapeutic stem cell preparation exert immunomodulatory effects through release of phosphatidylserine. Signal Transduct Target Ther. 2021 Jul 14;6(1):270.

13. Kholodenko IV, Kholodenko RV, Majouga AG, Yarygin KN. Apoptotic MSCs and MSC-Derived Apoptotic Bodies as New Therapeutic Tools. Curr Issues Mol Biol. 2022 Oct 24;44(11):5153–72.

14. Humbert P, Brennan MÁ, De Lima J, Brion R, Adrait A, Charrier C, et al. Apoptotic mesenchymal stromal cells support osteoclastogenesis while inhibiting multinucleated giant cells formation in vitro. Sci Rep. 2021 Jun 9;11(1):12144.

15. Vidula N, Villa M, Helenowski IB, Merchant M, Jovanovic BD, Meagher R, et al. Adverse Events During Hematopoietic Stem Cell Infusion: Analysis of the Infusion Product. Clin Lymphoma Myeloma Leuk. 2015 Nov;15(11):e157–162.

16. Musiał-Wysocka A, Kot M, Majka M. The Pros and Cons of Mesenchymal Stem Cell-Based Therapies. Cell Transplant. 2019 Jul;28(7):801–12.

17. Caruso S, Poon IKH. Apoptotic Cell-Derived Extracellular Vesicles: More Than Just Debris. Front Immunol. 2018 Jun 28;9:1486.

18. Chen H, Kasagi S, Chia C, Zhang D, Tu E, Wu R, et al. Extracellular Vesicles from Apoptotic Cells Promote TGFβ Production in Macrophages and Suppress Experimental Colitis. Sci Rep. 2019 Apr 10;9(1):5875.

19. Ma L, Chen C, Liu D, Huang Z, Li J, Liu H, et al. Apoptotic extracellular vesicles are metabolized regulators nurturing the skin and hair. Bioact Mater. 2022 May 11;19:626–41.

20. Wang J, Cao Z, Wang P, Zhang X, Tang J, He Y, et al. Apoptotic Extracellular Vesicles Ameliorate Multiple Myeloma by Restoring Fas-Mediated Apoptosis. ACS Nano. 2021 Sep 28;15(9):14360–72.

21. Zheng C, Sui B, Zhang X, Hu J, Chen J, Liu J, et al. Apoptotic vesicles restore liver macrophage homeostasis to counteract type 2 diabetes. J Extracell Vesicles. 2021 May;10(7):e12109.

22. Li M, Liao L, Tian W. Extracellular Vesicles Derived From Apoptotic Cells: An Essential Link Between Death and Regeneration. Front Cell Dev Biol [Internet]. 2020 [cited 2024 Jan 21];8. Available from:https://www.frontiersin.org/articles/10.3389/fcell.2020.573511

23. Liu PF, Hu YC, Kang BH, Tseng YK, Wu PC, Liang CC, et al. Expression levels of cleaved caspase-3 and caspase-3 in tumorigenesis and prognosis of oral tongue squamous cell carcinoma. PLoS ONE. 2017 Jul 10;12(7):e0180620.

24. Théry C, Witwer KW, Aikawa E, Alcaraz MJ, Anderson JD, Andriantsitohaina R, et al. Minimal information for studies of extracellular vesicles 2018 (MISEV2018): a position statement of the International Society for Extracellular Vesicles and update of the MISEV2014 guidelines. J Extracell Vesicles. 2018 Dec 1;7(1):1535750.

25. Liang G, Kan S, Zhu Y, Feng S, Feng W, Gao S. Engineered exosome-mediated delivery of functionally active miR-26a and its enhanced suppression effect in HepG2 cells. Int J Nanomedicine. 2018 Jan 30;13:585–99.

26. Hill C, Dellar ER, Baena-Lopez LA. Caspases help to spread the message via extracellular vesicles. FEBS J. 2023 Apr;290(8):1954–72.

27. Watanabe S, Alexander M, Misharin AV, Budinger GRS. The role of macrophages in the resolution of inflammation. J Clin Invest. 2019 May 20;129(7):2619–28.

28. Shapouri-Moghaddam A, Mohammadian S, Vazini H, Taghadosi M, Esmaeili SA, Mardani F, et al. Macrophage plasticity, polarization, and function in health and disease. J Cell Physiol. 2018 Sep;233(9):6425–40.

29. Das A, Sinha M, Datta S, Abas M, Chaffee S, Sen CK, et al. Monocyte and macrophage plasticity in tissue repair and regeneration. Am J Pathol. 2015 Oct;185(10):2596–606.

30. Cui Y, Chen J, Zhang Z, Shi H, Sun W, Yi Q. The role of AMPK in macrophage metabolism, function and polarisation. J Transl Med. 2023 Dec 8;21(1):892.

31. Galipeau J. Macrophages at the nexus of mesenchymal stromal cell potency: The emerging role of chemokine cooperativity. Stem Cells Dayt Ohio. 2021 Sep;39(9):1145–54.

32. Zhu Y, Zhang X, Yang K, Shao Y, Gu R, Liu X, et al. Macrophage-derived apoptotic vesicles regulate fate commitment of mesenchymal stem cells via miR155. Stem Cell Res Ther. 2022 Jul 16;13(1):323.

33. Ko JH, Kim HJ, Jeong HJ, Lee HJ, Oh JY. Mesenchymal Stem and Stromal Cells Harness Macrophage-Derived Amphiregulin to Maintain Tissue Homeostasis. Cell Rep. 2020 Mar 17;30(11):3806–3820.e6.

34. Liang ZY, Xu XJ, Rao J, Yang ZL, Wang CH, Chen CM. Mesenchymal Stem Cell-Derived Exosomal MiRNAs Promote M2 Macrophages Polarization: Therapeutic Opportunities for Spinal Cord Injury. Front Mol Neurosci. 2022 Jul 12;15:926928.

35. Patil M, Saheera S, Dubey PK, Kahn-Krell A, Kumar Govindappa P, Singh S, et al. Novel Mechanisms of Exosome-Mediated Phagocytosis of Dead Cells in Injured Heart. Circ Res. 2021 Nov 12;129(11):1006–20.

36. Fujii S, Miura Y, Fujishiro A, Shindo T, Shimazu Y, Hirai H, et al. Graft-Versus-Host Disease Amelioration by Human Bone Marrow Mesenchymal Stromal/Stem Cell-Derived Extracellular Vesicles Is Associated with Peripheral Preservation of Naive T Cell Populations. Stem Cells Dayt Ohio. 2018 Mar;36(3):434–45.

37. Lai P, Chen X, Guo L, Wang Y, Liu X, Liu Y, et al. A potent immunomodulatory role of exosomes derived from mesenchymal stromal cells in preventing cGVHD. J Hematol OncolJ Hematol Oncol. 2018 Dec 7;11(1):135.

38. Zhao RZ, Jiang S, Zhang L, Yu ZB. Mitochondrial electron transport chain, ROS generation and uncoupling (Review). Int J Mol Med. 2019 Jul;44(1):3–15.

39. Hernansanz-Agustín P, Enríquez JA. Generation of Reactive Oxygen Species by Mitochondria. Antioxid Basel Switz. 2021 Mar 9;10(3):415.

40. Gupta S, Pinky null, Vishal null, Sharma H, Soni N, Rao EP, et al. Comparative Evaluation of Anti-Fibrotic Effect of Tissue Specific Mesenchymal Stem Cells Derived Extracellular Vesicles for the Amelioration of CCl4 Induced Chronic Liver Injury. Stem Cell Rev Rep. 2022 Mar;18(3):1097–112.

41. Luo Q, Xian P, Wang T, Wu S, Sun T, Wang W, et al. Antioxidant activity of mesenchymal stem cell-derived extracellular vesicles restores hippocampal neurons following seizure damage. Theranostics. 2021;11(12):5986–6005.

42. Xiao X, Xu M, Yu H, Wang L, Li X, Rak J, et al. Mesenchymal stem cell-derived small extracellular vesicles mitigate oxidative stress-induced senescence in endothelial cells via regulation of miR-146a/Src. Signal Transduct Target Ther. 2021 Oct 22;6(1):1–15.

43. Hashim Z, Ilyas A, Zarina S. Therapeutic effect of hydrogen peroxide via altered expression of glutathione S-transferase and peroxiredoxin-2 in hepatocellular carcinoma. Hepatobiliary Pancreat Dis Int. 2020 Jun 1;19(3):258–65.

44. Tseng TH, Wang CJ, Lee YJ, Shao YC, Shen CH, Lee KC, et al. Suppression of the Proliferation of Huh7 Hepatoma Cells Involving the Downregulation of Mutant p53 Protein and Inactivation of the STAT 3 Pathway with Ailanthoidol. Int J Mol Sci. 2022 May 4;23(9):5102.

45. Li Y chao, Zheng J, Wang X zi, Wang X, Liu W jing, Gao J lu. Mesenchymal stem cell-derived exosomes protect trabecular meshwork from oxidative stress. Sci Rep. 2021 Jul 21;11(1):14863.

46. Damania A, Jaiman D, Teotia AK, Kumar A. Mesenchymal stromal cell-derived exosome-rich fractionated secretome confers a hepatoprotective effect in liver injury. Stem Cell Res Ther. 2018 Feb 6;9(1):31.

47. Paliwal S, Chaudhuri R, Agrawal A, Mohanty S. Human tissue-specific MSCs demonstrate differential mitochondria transfer abilities that may determine their regenerative abilities. Stem Cell Res Ther. 2018 Nov 8;9(1):298.

48. Maheshwari D, Kumar D, Jagdish RK, Nautiyal N, Hidam A, Kumari R, et al. Bioenergetic Failure Drives Functional Exhaustion of Monocytes in Acute-on-Chronic Liver Failure. Front Immunol. 2022;13:856587.

49. Faria J, Calcat-i-Cervera S, Skovronova R, Broeksma BC, Berends AJ, Zaal EA, et al. Mesenchymal stromal cells secretome restores bioenergetic and redox homeostasis in human proximal tubule cells after ischemic injury. Stem Cell Res Ther. 2023 Dec 10;14(1):353.

50. Phinney DG, Di Giuseppe M, Njah J, Sala E, Shiva S, St Croix CM, et al. Mesenchymal stem cells use extracellular vesicles to outsource mitophagy and shuttle microRNAs. Nat Commun. 2015 Oct 7;6(1):8472.

51. Morrison TJ, Jackson MV, Cunningham EK, Kissenpfennig A, McAuley DF, O’Kane CM, et al. Mesenchymal Stromal Cells Modulate Macrophages in Clinically Relevant Lung Injury Models by Extracellular Vesicle Mitochondrial Transfer. Am J Respir Crit Care Med. 2017 Nov 15;196(10):1275–86.

52. Russo E, Lee JY, Nguyen H, Corrao S, Anzalone R, La Rocca G, et al. Energy Metabolism Analysis of Three Different Mesenchymal Stem Cell Populations of Umbilical Cord Under Normal and Pathologic Conditions. Stem Cell Rev Rep. 2020 Jun;16(3):585–95.

53. Zhao R, Chen X, Song H, Bie Q, Zhang B. Dual Role of MSC-Derived Exosomes in Tumor Development. Stem Cells Int. 2020 Sep 9;2020:8844730.

54. Vallabhaneni KC, Penfornis P, Dhule S, Guillonneau F, Adams KV, Mo YY, et al. Extracellular vesicles from bone marrow mesenchymal stem/stromal cells transport tumor regulatory microRNA, proteins, and metabolites. Oncotarget. 2015 Mar 10;6(7):4953–67.

55. Lee HK, Finniss S, Cazacu S, Bucris E, Ziv-Av A, Xiang C, et al. Mesenchymal stem cells deliver synthetic microRNA mimics to glioma cells and glioma stem cells and inhibit their cell migration and self-renewal. Oncotarget. 2013 Feb;4(2):346–61.

56. Yue X, Gu M, Jia T. Upregulated miR-125b mitigates inflammation, astrocyte activation, and dysfunction of spinal cord injury by inactivating the MAPK pathway. Histol Histopathol. 2023 Jan 18;18624.

57. He M, Wu N, Leong MC, Zhang W, Ye Z, Li R, et al. miR-145 improves metabolic inflammatory disease through multiple pathways. J Mol Cell Biol. 2019 Apr 3;12(2):152–62.

58. Li HW, Zeng HS. Regulation of JAK/STAT signal pathway by miR-21 in the pathogenesis of juvenile idiopathic arthritis. World J Pediatr WJP. 2020 Oct;16(5):502–13.

59. Song C, Liu B, Ge X, Li H, Liu B, Xu P. miR-34a/Notch1b mediated autophagy and apoptosis contributes to oxidative stress amelioration by emodin in the intestine of teleost Megalobrama amblycephala. Aquaculture. 2022 Jan 30;547:737441.

60. Wang Y, Yin Z, Zhang N, Song H, Zhang Q, Hao X, et al. MiR-125a-3p inhibits cell proliferation and inflammation responses in fibroblast-like synovial cells in rheumatoid arthritis by mediating the Wnt/β-catenin and NF-κB pathways via targeting MAST3. J Musculoskelet Neuronal Interact. 2021;21(4):560–7.

61. Mei Y, Bian C, Li J, Du Z, Zhou H, Yang Z, et al. miR-21 modulates the ERK-MAPK signaling pathway by regulating SPRY2 expression during human mesenchymal stem cell differentiation. J Cell Biochem. 2013 Jun;114(6):1374–84.

62. Hart M, Walch-Rückheim B, Friedmann KS, Rheinheimer S, Tänzer T, Glombitza B, et al. miR-34a: a new player in the regulation of T cell function by modulation of NF-κB signaling. Cell Death Dis. 2019 Jan 18;10(2):1–14.

63. Liao W, He XJ, Zhang W, Chen YL, Yang J, Xiang W, et al. MiR-145 participates in the development of lupus nephritis by targeting CSF1 to regulate the JAK/STAT signaling pathway. Cytokine. 2022 Jun;154:155877.

64. Gupta S, Rawat S, Arora V, Kottarath SK, Dinda AK, Vaishnav PK, et al. An improvised one-step sucrose cushion ultracentrifugation method for exosome isolation from culture supernatants of mesenchymal stem cells. Stem Cell Res Ther. 2018 Jul 4;9(1):180.

65. Abuawad A, Mbadugha C, Ghaemmaghami AM, Kim DH. Metabolic characterisation of THP-1 macrophage polarisation using LC–MS-based metabolite profiling. Metabolomics. 2020;16(3):33.

66. Margiana R, Markov A, Zekiy AO, Hamza MU, Al-Dabbagh KA, Al-Zubaidi SH, Hameed NM, Ahmad I, Sivaraman R, Kzar HH, Al-Gazally ME. Clinical application of mesenchymal stem cell in regenerative medicine: a narrative review. Stem Cell Research & Therapy. 2022 Dec;13(1):1–22.

67. Han Y, Yang J, Fang J, Zhou Y, Candi E, Wang J, Hua D, Shao C, Shi Y. The secretion profile of mesenchymal stem cells and potential applications in treating human diseases. Signal Transduction and Targeted Therapy. 2022 Mar 21;7(1):92.

68. Banerjee P, Gaddam N, Chandler V, Chakraborty S. Oxidative Stress-induced liver damage and remodeling of the liver vasculature. The American Journal of Pathology. 2023 Jun 23.

